# A Haplotype-resolved Telomere-to-Telomere Pig Genome

**DOI:** 10.64898/2026.04.02.716236

**Authors:** Chengchen Zhao, Zheting Zhang, Zejie Lin, Jiajun Li, Wei Shi, Yuhao Wu, MaoXiang Shi, Tianci Kong, Bo Wang, Bingbo Shi, Xiaomin Wang, Jinzhu Xiang, Changjiang Xu, Yu Fu, Jin Ming, Yue Qin, Junqi Kuang, Hanning Wang, Yuxiang Yao, Bo Wang, Duanqing Pei

## Abstract

Haplotype-resolved telomere-to-telomere (T2T) pig genomes have not been reported, but may be essential for xenotransplantation and interspecies chimerism. Here we describe fully phased paternal and maternal T2T assemblies of an Erhualian pig (*Ssc_EHL*), enabling chromosome-scale reconstruction at base resolution. We uncover 2,498 (paternal haplotype) and 2,577 (maternal haplotype) protein-coding loci previously unresolved in *Sscrofa11*.*1*, while maintaining high concordance for anchored gene models between haplotypes. Applying this haplotype-resolved T2T annotation to publicly available pig-to-human kidney xenograft transcriptomes further allows the resolution of xenograft cell states and reveals newly resolved, cell-type-enriched genes. Cross-species synteny and orthology analyses against human identified a subset of high-confidence pig-specific genes with limited human counterparts, thus, refining the genomic basis of interspecies divergence. By integrating these orthology features with organ transcriptomes, we further defined an Interspecies Chimerism Compatibility score to prioritize organ-enriched, highly conserved loci with minimal redundancy imbalance, providing haplotype consistent targets for engineering organ-deficient pigs. Together, this haplotype-resolved T2T pig genome provides a foundational resource for mechanistic studies and rational genome engineering in xenotransplantation and interspecies chimerism.

## Introduction

Pigs (Sus scrofa domesticus) are omnivores and have been domesticated along with chicken and dogs, figuring quite prominently in human society, especially in Asia. Over 1.5 billion of pigs are raised annually to provide food to the human population worldwide. Pig breeding has becoming an important means to boost pig farming efficiency and improve quality for human health. Remarkably, pig organs such as liver, kidney and heart have been transplanted into human to sustain life, marking a new era in medical history^1–5^. The breakthrough in pig xenotransplantation is made possible through extensive genetic modification of the pig genome, with more than 10 distinct edits. It is conceivable that future improvements can be made through more editing such that the organ can survive the insult from the immune system of the recipient and better adaptation of the organ to the host.

Unfortunately, the pig genome remains poorly characterized, compared to the completion of high quality T2T genomes from human^6^, mouse^7^, and macaque^8^. One attempt yielded only near-T2T assemblies, including the Jinhua genome^9^ (six remaining gaps), Rongchang genome and Min genome^10^ (lacking chr Y), alongside a hybrid Duroc genome combining a female Duroc genome with a separate Y chromosome. Most recently, a T2T assembly of the Wuzhishan pig^11^ (T2T-pig1.0) provided an important advance, but its haplotypes still contain 51 paternal and 38 maternal gaps and lack definitive parental phasing due to the absence of parental sequencing data.

We report here a haplotype-resolved T2T genome for the Erhualian pig (*Ssc_EHL*), assembled using PacBio HiFi, Oxford Nanopore ultra-long, MGI short-read, and Pore-C sequencing data. The assembly achieves complete phasing, yielding gap-free paternal and maternal haplotypes for all chromosomes, including both X and Y. This resource establishes a definitive reference standard for porcine genomics and provides a foundation for dissecting allelic diversity, structural heterozygosity, and the genomic determinants of developmental competence. Beyond resolving novel centromere patterns, the *Ssc_EHL* T2T genome uncovers numerous pig-specific genes that may be candidates as targets for potential genome editing to improve xenotransplantation. Furthermore, we attempted to curate a candidate list of targets for generating humanized organs through interspecies chimerism.

## Results

### Haplotype-resolved T2T Erhualian Genome

We have been interested in pig as a model organism and have generated the first porcine iPSCs in 2010^12^ and porcine blastoids in 2024^13^. In recent years, we have become more interested in the Erhualian pig (*Ssc_EHL*) which is a local breed in eastern China known for its large sizes of piglings per pregnancy (up to 40). To further develop it into a model organism for regenerative medicine, we set out to establish a haplotype-resolved T2T reference genome for the Erhualian pig.

We intergrated high-coverage PacBio HiFi (198.51 Gbp, 76.35×), Oxford Nanopore ultra-long reads (268.33 Gbp, 103.20×), and MGI short reads (126.45 Gbp, 48.64×), with Pore-C facilitating chromosomal scaffolding (101.34 Gbp, 38.98×) (Supplementary Table 1) while leveraging a trio-binning strategy supported by parental sequencing data, to obtain full phased, gap-free paternal (*Ssc_EHL* haplotype1, hap1) and maternal (*Ssc_EHL* haplotype2, hap2) assemblies at the chromosome level (Supplementary Tables 2, 3). The final assemblies achieved unprecedented contiguity, with contig N50 values of 152.92 Mb (paternal) and 153.50 Mb (maternal). This contiguity substantially exceeds all prior assemblies. For comparison, the Jinhua assembly (JH-T2T, labeled as *Ssc_JH* in the following text) retains 116 maternal and 42 paternal gaps after gap-closing efforts. The Rongchang (T2T_RCpig1.0, *Ssc_RC*) and Min (T2T_Mpig1.0, *Ssc_MIN*) assemblies do not include the Y chromosome. The Wuzhishan assembly (T2T-pig1.0, *Ssc_WZS*) rely on non-parental phasing methods that leave 51 maternal and 38 paternal gaps. Whole-genome alignments to *Sscrofa11*.*1* showed broad synteny, with difference concentrated in discrete inversions, translocations, and duplication blocks (Fig. 1a).

**Fig. 1.**
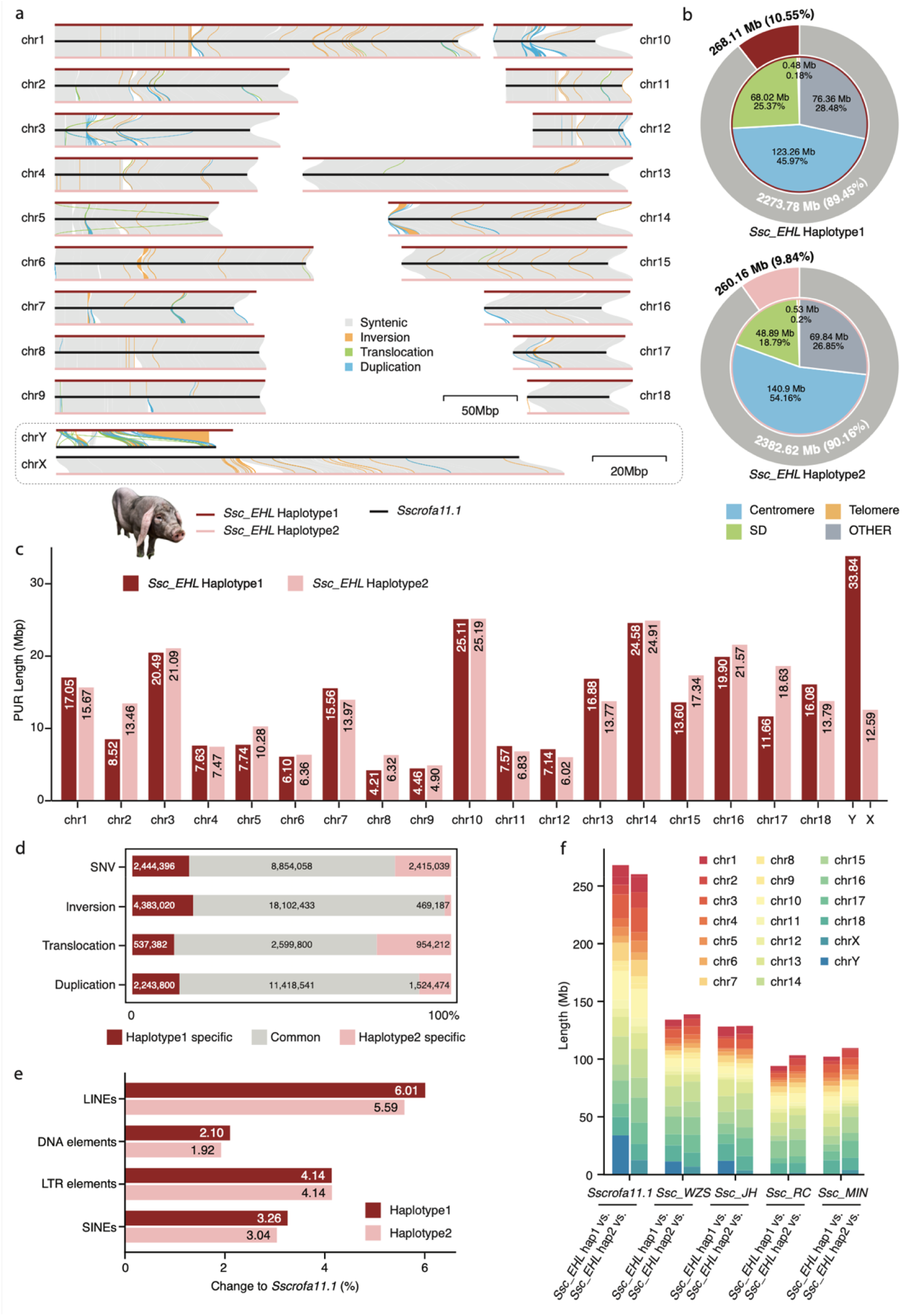
Overview of the haplotype-resolved T2T Erhualian pig genome and structural diversity. **(a)** Whole-genome alignments of Ssc_EHL hap1 (paternal, dark red) and hap2 (maternal, light red) against the Sscrofa11.1 reference (black). Colored links represent syntenic blocks and major structural differences, including inversions, translocations, and duplications. Scale bars: 50 Mb (autosomes), 20 Mb (sex chromosomes). **(b)** Recovery of previously unresolved regions (PURs) in Ssc_EHL assemblies relative to Sscrofa11.1. Outer rings indicate the fraction of PUR versus non-PUR sequences per haplotype; inner pie charts partition PURs into centromeric, segmental duplication (SD), telomeric, and other categories. **(c)** Chromosome-wide PUR lengths (Mb) for Ssc_EHL hap1 and hap2. **(d)** Partitioning of intra-individual variation into hap1-specific, shared, and hap2-specific components across SNV and SVs. **(e)** Divergence in major repeat classes between Ssc_EHL haplotypes and Sscrofa11.1, expressed as percent change relative to the reference. **(f)** Cross-breed comparison of PURs using Ssc_EHL haplotypes as references against Sscrofa11.1 and near-T2T assemblies (Ssc_WZS, Ssc_JH, Ssc_RC, and Ssc_MIN). Bar heights represent total PUR length, with colors indicating chromosomal contributions.

The paternal and maternal assemblies span 2.54 Gb and 2.64 Gb, recovering 268.11 Mb and 260.16 Mb of previously unresolved sequence relative to *Sscrofa11*.*1*, respectively. These previously unresolved regions (PURs) are predominantly composed of structurally complex elements, including centromeric satellites, segmental duplications, and telomeric arrays (Fig. 1b, Supplementary Table 4). The centromeric regions are dominated by tandem satellite arrays with a predominant ~335 bp monomer and support by overlapping CENP-A CUT&RUN and published ChIP-seq^11^ signals (Supplementary Fig. 1a-c). Telomeric tracts are present at both 5′ and 3′ chromosome ends across haplotypes, supporting end-to-end telomere representation in the T2T assemblies (Supplementary Fig. 1d). The assembly quality is quite high, with BUSCO completeness of 99.89% for hap2 and 97.67% for hap1. These BUSCO values exceed or are comparable to those reported for recent T2T assemblies, including Jinhua (*Ssc_JH*, 96.4%), Min (*Ssc_MIN*, 98.2%), Rongchang (*Ssc_RC*, 98.3%), and Wuzhishan (*Ssc_WZS*, 96.2%). Merqury analysis yielded consensus quality values (QV) exceeding 50 (>99.999% accuracy) for autosomes in both haplotype and X chromosome, with the Y chromosome also achieving a high-fidelity QV of 45.8 (Supplementary Table 5). These metrics further reveal the robust construction of a contiguous, gap-free physical map, verifying the integrity of assembly across previously unsolved genomic regions. Quantitative analysis of PURs reveals distinct landscapes across chromosomes, highlighting structural divergence between paternal and maternal haplotypes (Fig. 1c).

Leveraging the fully phased assembly, we profiled haplotype divergence between the paternal and maternal haplotypes at both single nucleotide variant (SNV) and structural variant (SV) scales. Genome-wide SNV density averaged 4.35 per kb, with low-variation tracts interspersed with hotspots of elevated divergence. Across variant types, most variant burden is shared between haplotypes, whereas the haplotype-specific fraction differed by category (Fig. 1d). Inversions and duplications were more frequently observed in hap1, whereas translocations were more frequently observed in hap2. Repeat annotation further showed that repetitive sequences account for 45.33% of the paternal haplotype and 45.50% of the maternal haplotype. While major repeat classes are broadly similar between haplotypes, we observed marked divergence in LINEs and SINEs, underscoring the rapid evolutionary turnover of these structurally complex regions (Table 1, Fig. 1e). Consistent with this pattern, segmental duplications (SD) are concentrated in centromeric and pericentromeric regions across both haplotypes and are largely shared, with a smaller haplotype-specific fraction enriched on sex chromosomes (Supplementary Fig. 2a-c, Supplementary Table 6). Many SD intervals overlap with annotated genes and are enriched for innate immune and antiviral response programs, indicating that structurally complex regions contribute not only to copy-number diversification but also to functional variation between haplotypes (Supplementary Fig. 2d). Such variation in repeat-rich regions has practical implications, particularly for porcine endogenous retroviruses (PERVs), whose accurate identification and mapping for inactivation are highly sensitive to reference completeness^14^. Using the *Ssc_EHL* assembly, we identified a large number of PERV loci with a nonrandom chromosomal distribution, many of which mapped within PURs (Supplementary Fig. 3a).

**Table 1.**
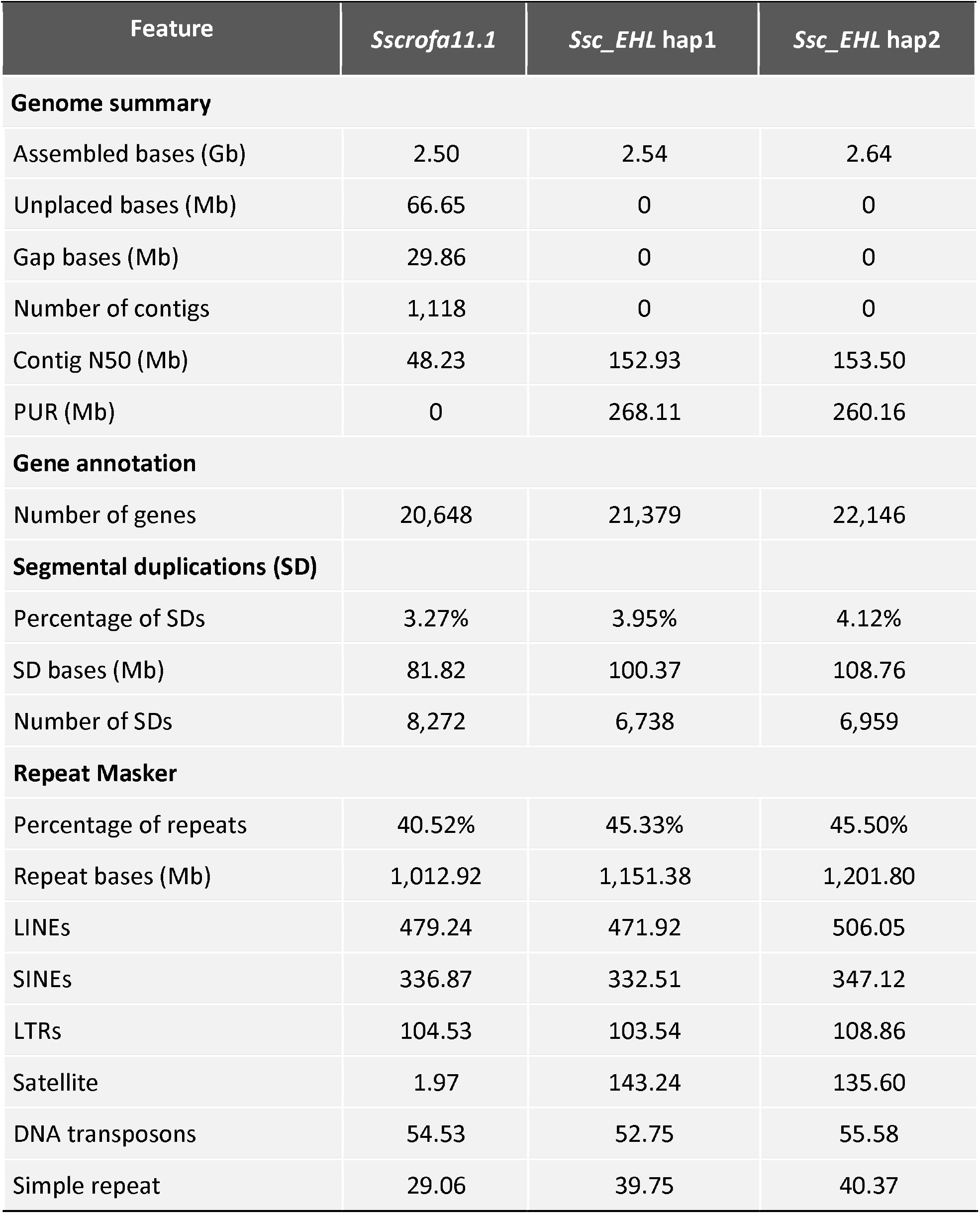
Comparison of pig genome assemblies between *Sscrofa11.1* and *Ssc_EHL*.

| Feature | *Sscrofa11.1* | *Ssc_EHL* hap1 | *Ssc_EHL* hap2 |
| --- | --- | --- | --- |
| Genome summary |  |  |  |
| Assembled bases (Gb) | 2.50 | 2.54 | 2.64 |
| Unplaced bases (Mb) | 66.65 | 0 | 0 |
| Gap bases (Mb) | 29.86 | 0 | 0 |
| Number of contigs | 1,118 | 0 | 0 |
| Contig N50 (Mb) | 48.23 | 152.93 | 153.50 |
| PUR (Mb) | 0 | 268.11 | 260.16 |
| **Gene annotation** |  |  |  |
| Number of genes | 20,648 | 21,379 | 22,146 |
| **Segmental duplications (SD)** |  |  |  |
| Percentage of SDs | 3.27% | 3.95% | 4.12% |
| SD bases (Mb) | 81.82 | 100.37 | 108.76 |
| Number of SDs | 8,272 | 6,738 | 6,959 |
| **Repeat Masker** |  |  |  |
| Percentage of repeats | 40.52% | 45.33% | 45.50% |
| Repeat bases (Mb) | 1,012.92 | 1,151.38 | 1,201.80 |
| LINEs | 479.24 | 471.92 | 506.05 |
| SINEs | 336.87 | 332.51 | 347.12 |
| LTRs | 104.53 | 103.54 | 108.86 |
| Satellite | 1.97 | 143.24 | 135.60 |
| DNA transposons | 54.53 | 52.75 | 55.58 |
| Simple repeat | 29.06 | 39.75 | 40.37 |

In contrast to the modest intra-individual differences observed between the two *Ssc_EHL* haplotypes, inter-breed comparisons reveal substantially greater divergence in PUR content, SV, and repeat composition. Across published pig genomes, PURs are broadly conserved in composition but differ markedly in total length and chromosomal distribution, consistent with lineage-specific expansion of repetitive and duplicated sequences. Using both *Ssc_EHL* haplotypes as references, we observed concordant PUR hotspots between haplotypes, with recurrently localized to pericentromeric and subtelomeric regions across breeds (Supplementary Fig. 3b, c). Beyond these shared hotspots, additional PURs are also present in interstitial regions in a breed-dependent manner, indicating lineage-specific structural diversification outside centromeres (Supplementary Fig. 4). To separate the contribution of centromeric assemblies from other sequences, we further quantified PUR lengths after excluding centromeric regions. Substantial non-centromeric PURs remain across genomes, revealing pronounced between-breed differences beyond centromeric assembly status (Fig. 1f, Supplementary Fig. 3d). We next systematically quantified SV across these genomes. Although more than 2.25 Gb of each genome aligned within syntenic blocks, the scale and composition of rearrangements differ markedly among breeds (Supplementary Table 7, Supplementary Fig. 5). In particular, duplication burden varies substantially, with *Ssc_WZS* and *Ssc_JH* exhibiting the largest duplication length, whereas *Ssc_RC* and *Ssc_MIN* show comparatively lower duplication and inversion burden. These patterns are consistent across both *Ssc_EHL* haplotypes, supporting lineage-specific expansion of duplicated regions rather than haplotype-specific effects. Repeat content also varies among breeds in both genomic fraction and absolute annotated bases across major repeat classes (LINEs, SINEs, LTRs, satellites, DNA transposons, simple repeats, and low-complexity sequences), consistent with lineage-specific repeat expansion and turnover (Supplementary Table 8). Collectively, these results demonstrate that inter-breed structural heterogeneity substantially exceeds intra-individual haplotype differences, and that much of this divergence arises from lineage-specific diversification in duplicated and rearranged regions that are largely inaccessible without T2T assemblies. Based on these characterization, we conclude that the Erhualian T2T genome provides the first complete diploid sequence information of pig.

### Novel Pig Genes from Paternal and Maternal Haplotypes

The complete sequences provided by T2T Erhualian assemblies offer a unique opportunity to annotate novel genes. We integrated ab initio predictions with evidence-based gene models from transcriptome data, reference-based projection (Liftoff) and protein-to-genome alignment (miniprot) to annotate each haplotype. Gene models from the draft T2T annotation, Liftoff, and miniprot have been reconciled locus-by-locus, and the best-supported model is retained. Loci supported by Liftoff or miniprot but absent from the draft set are kept as “Unanchored”, whereas well-supported T2T-only loci are retained as “Novel” (Supplementary Fig. 6).

In total, we annotated 21,379 or 22,146 protein-coding genes, including 2,498 or 2,577 novel unresolved in *Sscrofa11*.*1* on hap1 (paternal) or hap2 (maternal), respectively (Fig. 2a). We also annotated 6,471 ncRNAs in hap1 and 6,741 in hap2 (Supplementary Table 9). Among these additional protein-coding genes, 2,068 in hap1 and 2,116 in hap2 are supported by both RNA-seq evidence and functional annotation. In addition, 333 genes in hap1 and 347 in hap2 are supported by functional annotation only, whereas 97 genes in hap1 and 112 in hap2 are supported by RNA-seq only (Fig. 2b). Comparison of gene models between haplotypes shows high concordance, with 18,971 genes anchored in syntenic positions and only a small fraction classified as relocated (52) or multi-hit (201 in hap1 and 193 in hap2) (Fig. 2c). The remaining unanchored fractions (2,133 in hap1 and 2,909 in hap2) are largely attributable to loci lacking one-to-one counterparts between haplotypes, consistent with sex-chromosome-specific content. Among these unanchored genes, 1,406 in hap1 and 1,454 in hap2 retain protein-level homology to the other one, based on best cross-haplotype protein BLAST hits meeting the thresholds of bitscore ≥ 200 and identity ≥ 80%. The haplotype-resolved annotations provide a consistent gene catalog across parental haplotypes while retaining haplotype-specific loci that would be collapsed or missed in a single-reference framework.

**Fig. 2.**
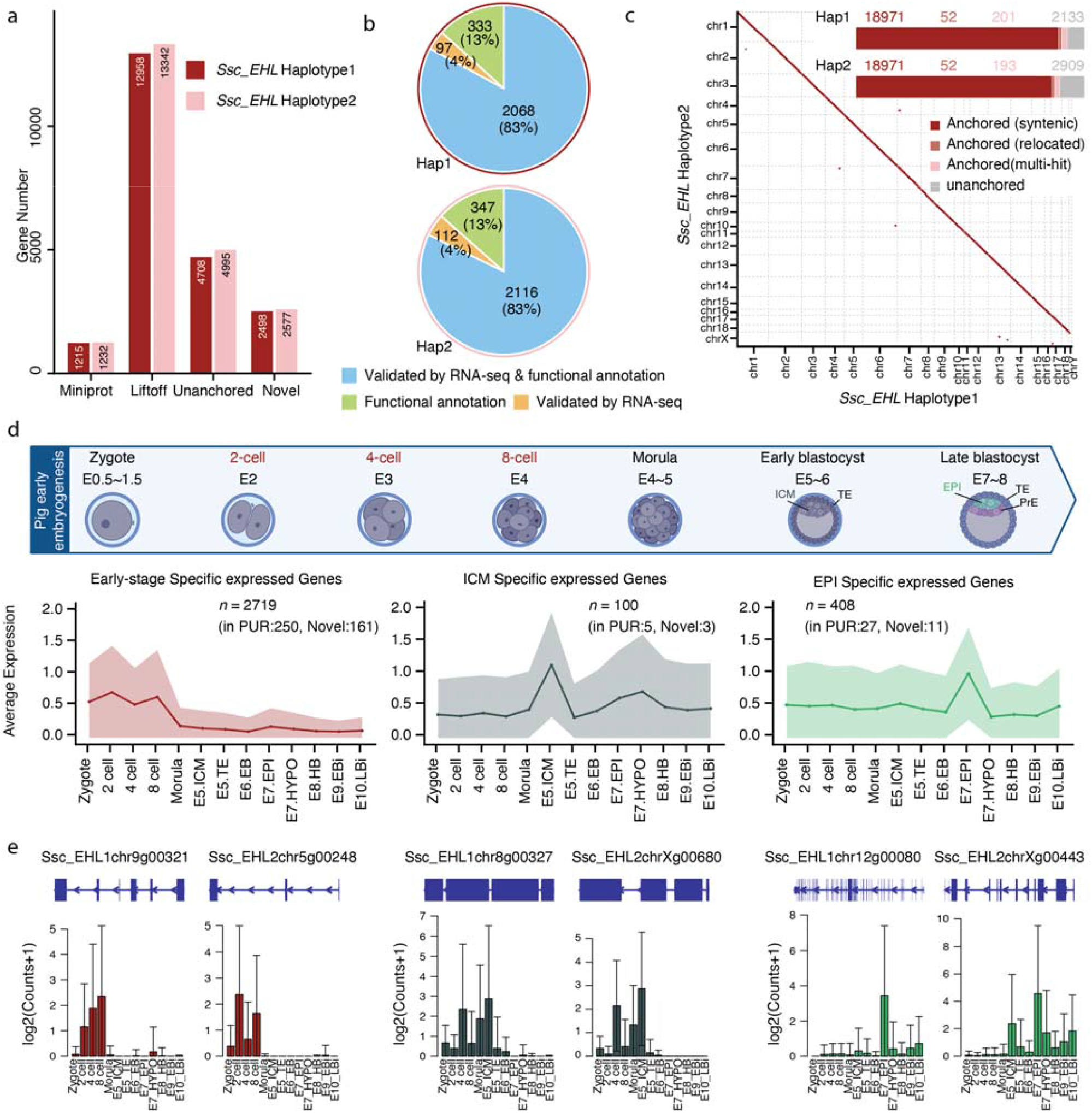
Gene annotation of the T2T assemblies and developmental relevance during early embryogenesis. **(a)** Counts of protein-coding gene models for Ssc_EHL hap1 and hap2, categorized by source: Miniprot-derived, Liftoff-projected, Unanchored loci, and T2T-only Novel loci. **(b)** Evidence supporting additional protein-coding genes, classified by RNA-seq validation and/or functional annotation. **(c)** Haplotype-to-haplotype comparison of gene model positions, summarizing concordance based on syntenic anchoring, relocation, or multi-hit mappings (red points). **(d)** Average expression profiles for early-stage specific genes, ICM-specific genes, and epiblast-specific genes. **(e)** Representative examples of stage specific genes with curated gene models and their corresponding expression profiles.

Given the substantially large number of genes relative to previously annotated one, we investigated them further and show that they arise from multiple sources, including reannotation of previously unplaced scaffolds and improved resolution of duplicated regions (Supplementary Table 10). Among them, 124 genes in hap1 and 129 in hap2 correspond to reannotated genes that had previously resided on unplaced scaffolds, including functionally relevant loci such as FUT9, SNRPN, SNURF, TNFAIP2, MARK3, AKT1, SNAPC4, PMPCA, INPP5E, SEC16A, and NOTCH1, indicating that our T2T genomes can recover biologically relevant genes that are fragmented or absent from the previous reference. Another subset of genes represent additional copies of previously annotated ones, consistent with improved resolution of duplicated regions (41 genes in hap1 and 30 in hap2). Another 27 genes in hap1 and 28 in hap2 are best matched to Swiss-Prot proteins from non-pig species, such as GPR61_HUMAN, GDE_CANLF, TIM10_RAT, VMP1_PONAB, ANKR2_HUMAN, and ERLN1_PONAB, rather than to curated porcine entries. This suggests that the corresponding loci retain clear protein-level homology but remain incompletely represented in current pig annotations. The remaining majority, 2,306 genes in hap1 and 2,390 in hap2, have no match in the reviewed Swiss-Prot database, suggesting that these loci are poorly represented in current curated annotations rather than simply missing canonical genes. To further characterize these genes, we searched them against the unreviewed TrEMBL database. Although a small subset of them have high-similarity matches to previously predicted porcine genes, most still share less than 50% sequence identity with their best matches, consistent with substantial sequence divergence from currently annotated pig proteins. We then performed domain annotation and further show that 31 genes in hap1 and 36 in hap2 are transcription factors, with POU, zinc finger, and homeodomain motifs, whereas 68 genes in hap1 and 76 in hap2 encode proteins with predicted signal peptides, suggesting potential roles in transcriptional regulation or intercellular communication. Functionally, these loci are not random: based on predicted Gene Ontology annotations, most of these loci encode proteins with catalytic or binding activities and are predominantly assigned to metabolic process, biological regulation, and developmental process categories (Supplementary Table 10). We further incorporated them into downstream comparative analyses and developmental transcriptome profiling as described below.

### Key Novel Genes for Porcine Early Embryogenesis

The lack of high quality genome data has so far hampered our understanding of porcine early embryogenesis and it remains problematic to develop and maintain stem cells with toti- and naïve-pluripotent potentials comparable to those reported for other species including mouse^15–17^ and human^18^. Thus, it is of great interest to see if the T2T genome with novel annotated genes may help to resolve this. To this end, we profiled gene expression across key stages and lineages of pig embryogenesis using public RNA-seq^19^ and single-cell RNA-seq^20^ (scRNA-seq) datasets (Supplementary Table 11), mapping reads to *Ssc_EHL* hap1 and hap2 separately. Based on temporal patterns, we defined three stage-specific gene sets: early-stages (2-cell to 8-cell stages, ESG, n=2,719), inner cell mass stage (ICM stage at early blastocyst, ISG, n=100), and epiblast stage (EPI stage at late blastocyst, EpSG, n=408) (Fig. 2d, Supplementary Table 12). Notably, 250 ESGs, 5 ISGs, and 27 EpSGs are located within PURs, among them, 161, 3, and 11, respectively, are novel genes (Fig. 2d). Previously reported totipotency-associated factors in mouse and human, including OBOX homologs (Ssc_EHL1chr6g00847 and Ssc_EHL1chr6g00849), ZSCAN4, and DUX homologs (DUXA, Ssc_EHL1chr10g00435, Ssc_EHL1chr10g00438, Ssc_EHL1chr10g00444, and Ssc_EHL1chr10g00448), are included in the ESG set (Supplementary Table 12, Supplementary Fig. 10). Representative novel genes have well-supported gene models and stage-restricted expression patterns on both haplotypes (Fig. 2e). In addition, 8 novel ESG genes in each haplotype encode predicted transcription factor domains, mostly homeodomains, and their predicted Gene Ontology terms are consistent with roles in regulatory programs. Genome-wide localization analysis further reveals their non-uniform chromosomal distribution with recurrent hotspots adjacent to centromeres and toward chromosome ends (Supplementary Fig. 7).

Together, these analyses demonstrate that haplotype-resolved T2T assemblies are essential for comprehensive annotation of genes from PUR-associated loci with stage- and lineage-restricted expression. These novel genes are high value candidates as key regulators of early embryonic programs, which have been absent from all previous assemblies.

### Comparative T2T Genomics for Pig vs Human or Mouse

We then compared the T2T genomes of Pig, mouse and Human. We first examined genomic divergence among pig, human, and mouse by integrating whole-genome synteny analysis with orthology-based homology inference. Whole-genome synteny analysis comparing *Ssc_EHL* with human and mouse reveal extensive conservation of chromosome-scale collinearity (Fig. 3a). At the genome scale, 92.2% and 91.0% of the *Ssc_EHL* genes are assigned to syntenic blocks relative to human and mouse, respectively (Fig.3b). While the proportion of pig genes falling within conserved syntenic regions exceeded 85% for most autosomes, this frequency dropped significantly on chr10 (66.3% vs. human, 71.6% vs. mouse) and chrY (25.9% vs. human, 24.3% vs. mouse) (Supplementary Fig. 8a). OrthoFinder-based orthogroup analysis shows that most genes are captured by cross-species orthogroups, with single-copy orthogroups representing the majority in pig-human (71.6%), pig-mouse (66.9%), and human-mouse (71.4%) comparisons (Fig. 3c). Protein sequence identity distributions for orthologous genes are strongly skewed toward high identity across all three pairwise comparisons, supporting high protein-level conservation among matched orthologs (Fig. 3d). Notably, the human-mouse comparison shows a single-copy orthogroup fraction (71.4%) comparable to pig-human (71.6%), whereas the protein-identity distributions broadly comparable. These results indicate that both species preserve extensive protein-level conservation to humans while differing in orthogroup structure and copy-number composition. Integrating orthogroup assignment and conserved synteny, 86.6% of pig genes are both orthogrouped and syntenic to human loci (Supplementary Fig. 8b), and developmental stage-specific gene sets show equal or higher concordance, consistent with broad conservation of core developmental programs (Supplementary Fig. 8b). Given the central role of pluripotency and totipotency regulators in early development, we further examined whether canonical stemness factors show comparable protein similarity across the three species. Averaged protein-level similarity scores indicated that pig-human similarity is higher than human-mouse for multiple representative factors, including SOX2, ESRRB, KLF4, MYC, and DUX (Fig. 3e, Supplementary Figs. 9, 10).

**Fig. 3.**
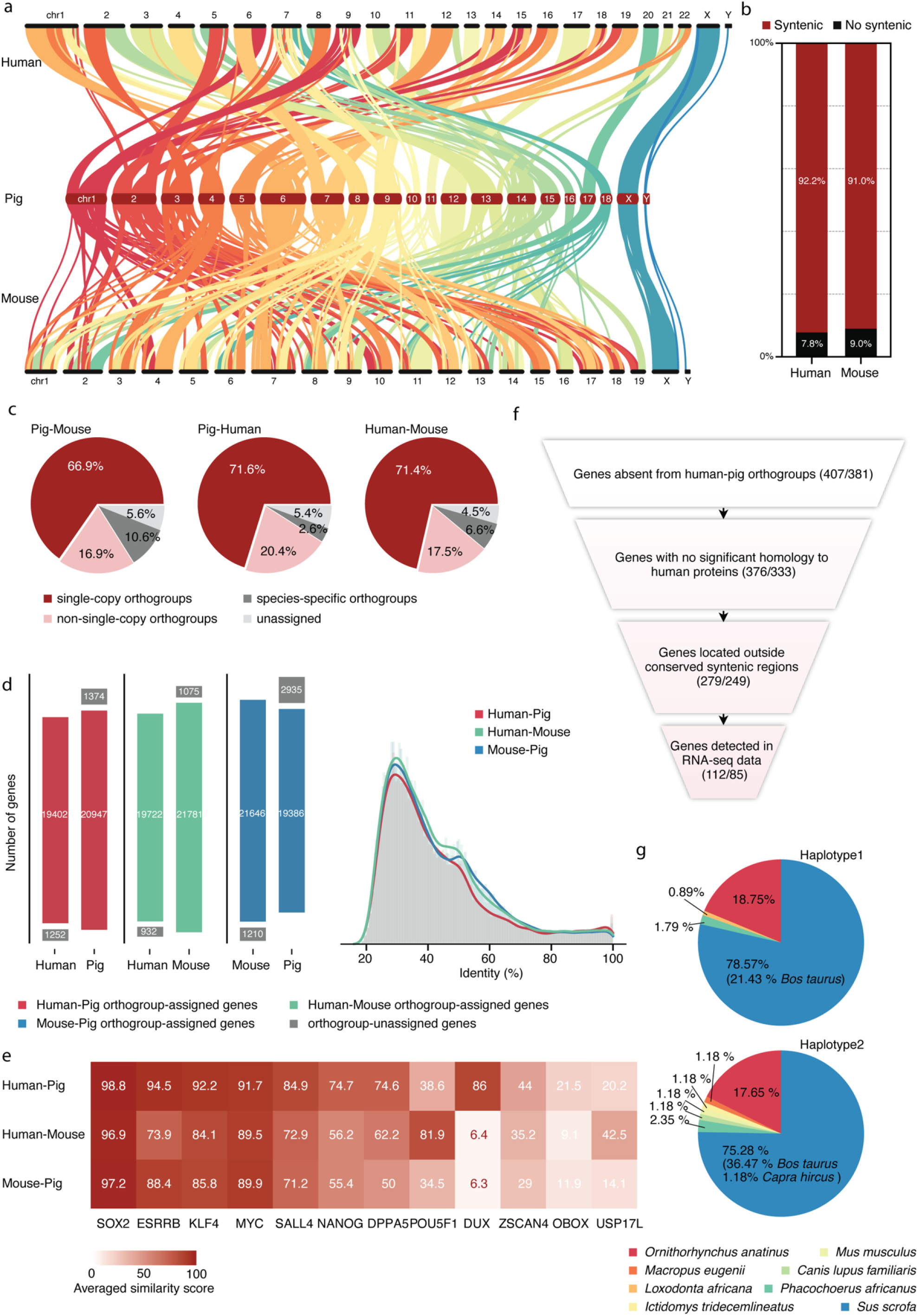
Cross-species synteny, orthology, and identification of pig-specific genes. **(a)** Chromosome-scale collinearity between human, pig (Ssc_EHL), and mouse genomes, ribbons connect conserved syntenic blocks. **(b)** Fraction of Ssc_EHL genes located within conserved syntenic blocks relative to human and mouse. **(c)** Orthogroup composition for pig-human and pig-mouse comparisons, partitioned into single-copy, non-single-copy, species-specific, and unassigned groups. **(d)** Barplots show counts of orthogy genes (left) and density plot shows protein sequence identity distributions (right) for pig-human and pig-mouse orthologous pairs. **(e)** Heatmap showing averaged protein similarity scores for selected pluripotency and totipotency factors defined in mouse in pairwise comparisons among human-pig, human-mouse, and mouse-pig. For each factor, the mouse protein was used as the query, and the closest matching ortholog(s) in the human and pig annotations, including both putative one-to-one orthologs and the most similar paralogs when necessary, were selected for similarity scoring. Color intensity reflects the averaged similarity score for each gene across the corresponding species pair. **(f)** Stepwise filtering strategy to define high-confidence pig-specific genes **(g)** Taxonomic distribution of top protein homology hits for pig-specific genes.

Despite this overall conservation, cross-species comparisons delineate a subset of expressed pig genes with limited human counterparts, motivating a focused search for high-confidence pig-specific candidates relevant to genetic risk assessment in xenotransplantation. We applied a stepwise filtering strategy (Supplementary Fig. 11a). Starting with genes absent from human-pig orthogroups, we removed genes with significant similarity to human proteins, excluded genes located within conserved syntenic regions, and yielded 279 candidate genes for hap1 and 249 for hap2. Among these, 112 genes in hap1 and 85 in hap2 are detected in our RNA-seq data (Supplementary Table 13, Fig. 3f). We next surveyed sequence similarity of these candidates across a representative panel of mammalian proteomes to contextualize their evolutionary origins (Supplementary Fig. 11b). Taxonomic assignment of top-scoring protein hits showed that most candidates match *Sus scrofa* entries, with only a minority mapping to other mammals, indicating limited detectable close homology outside pigs (Fig. 3g). Chromosomal mapping further revealed that these candidates are highly enriched within PURs, comprising 93.26% and 90.18% of genes for hap1 and hap2 (Supplementary Fig. 11c). Together, these analyses indicate that while genome-wide collinearity and most orthologous relationships are conserved between pig and human, a tractable set of pig-specific candidates persists outside conserved syntenic regions. These genes are preferentially associated with complex genomic contexts such as PURs, providing a focused set of loci for evaluating interspecies chimerism and organ transplantation compatibility as discussed below.

### Putative Genomic Determinants for Xenotransplantation

Pig kidneys have been transplanted into humans and apparently can sustain life for more than 217 days^4,21–25^, ushering in a new era of xenotransplantation and providing a viable alternative to human organs. Current success is predicated on gene editing to remove acute immune rejection of pig organ in human. However, the compatibility of pig organ to human physiology remains largely unresolved.

To further probe physiological compatibility beyond acute immune rejection, we reanalyzed single-cell transcriptomes from pig-to-human kidney xenografts^22^ using our T2T pig genome *Ssc_EHL*, which recovered the same 10 major renal cell populations reported previously, while resolving additional heterogeneity within the vimentin-positive proximal tubule (PT_VIM+) compartment, an injury-associated state linked to dedifferentiation, stress responses, and repair (Fig. 4a, Supplementary Fig. 12a). The improved reference enables finer stratification of this cell state and uncovers newly resolved, cell-type-enriched genes, exemplified by Ssc_EHL2chr1g02293 in endothelial cells, Ssc_EHL2chr17g00009 in PT_VIM+ cells, and Ssc_EHL1chr1g00164 in immune cells (Fig. 4b). For additional functional context beyond the primary gene annotation, we queried the predicted proteins against the unreviewed UniProtKB/TrEMBL database. The results suggest that Ssc_EHL2chr17g00009 corresponds to the small ribosomal subunit protein eS19, consistent with elevated translational programs in immune compartments and that Ssc_EHL2chr1g02293 corresponds to an EGFL7-like protein. Together, these newly resolved markers provide candidate molecular readouts of injury-associated proximal tubule states and xenograft cellular composition, and may refine assessment of pig kidney physiological compatibility in the human setting. To further characterize transcriptional responses during early xenograft adaptation, we reanalyzed the longitudinal bulk RNA-seq data generated from core biopsies collected at 0, 12, 24, and 48 hours (h) after transplantation using our T2T pig genome *Ssc_EHL*. Beyond the genes captured in the original reference, we show an additional set of previously unresolved loci with clear dynamic expression changes across time. Most of these genes are highly expressed at 0 h and rapidly decreased after transplantation, whereas a smaller subset shows delayed induction, particularly at 48 h. Further annotation indicates that these dynamically regulated genes include loci encoding predicted signal peptide-containing proteins and a transcription factor domain-containing gene, and those enriched for functions related to metabolic process, response to stimulus, and transmembrane transporter activity. Notably, genes with early loss of expression are mainly associated with intracellular metabolic and homeostatic functions, whereas genes induced at later time points more often with extracellular, signaling, or membrane-associated activities (Fig. 4c, Supplementary Fig. 12b). Together, these findings suggest that the our T2T assemblies would allow the uncovering of previously unknown transcriptional programs associated with the transition from baseline renal physiology to early xenograft stress and adaptation after pig-to-human kidney xenotransplantation.

**Fig. 4.**
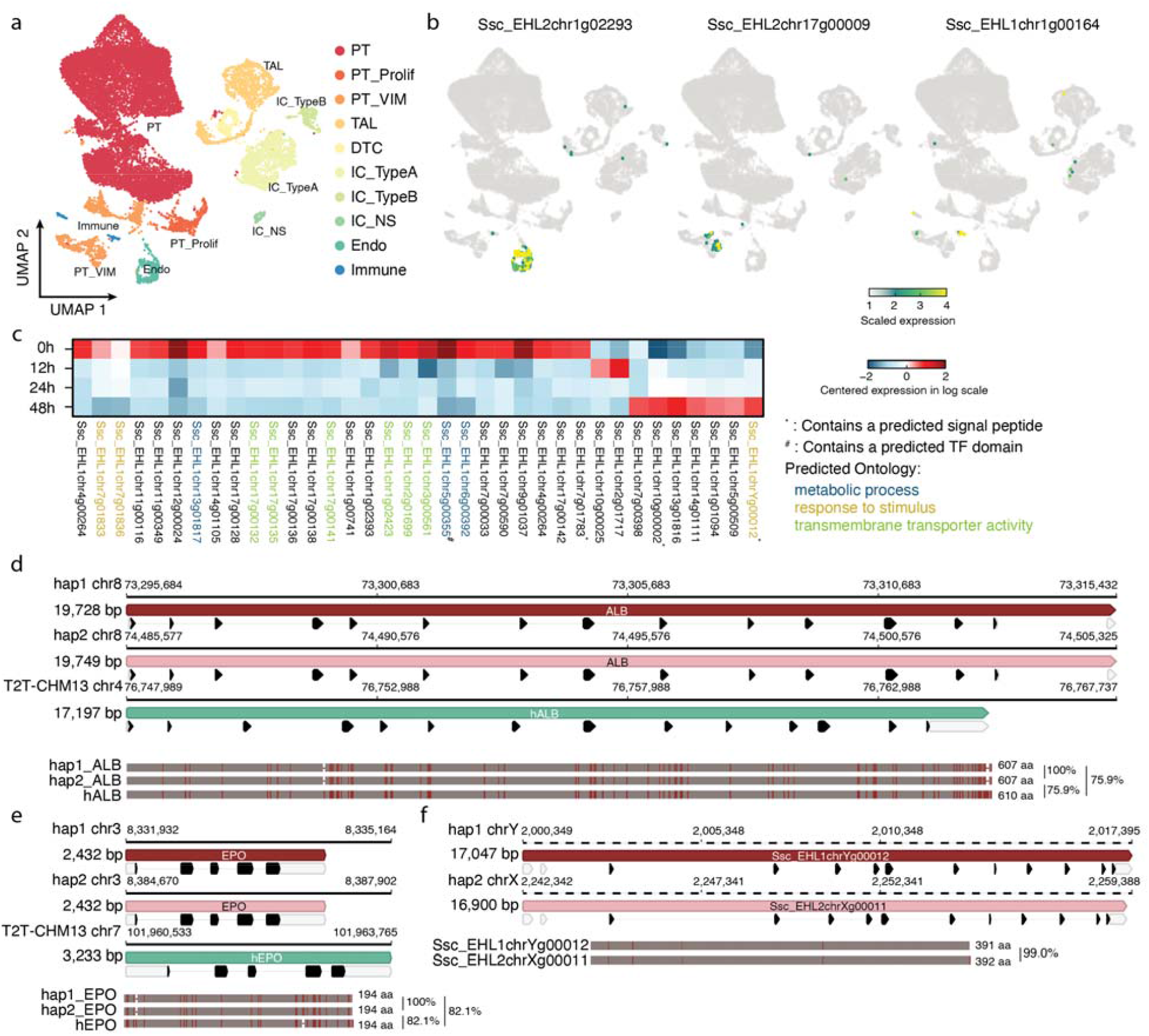
Genomic features relevant to xenotransplantation revealed by the T2T pig genome. **(a)** UMAP projection of single-cell transcriptomes from pig-to-human kidney xenografts, annotated into major populations including proximal tubule cells (PT), proliferating PT (PT_Prolif), vimentin-positive injury-associated PT (PT_VIM), thick ascending limb (TAL), distal tubule cells (DTC), type A and type B intercalated cells (IC_TypeA and IC_TypeB), nonspecific intercalated cells (IC_NS), endothelial cells (Endo), and immune cells. **(b)** Feature plots showing cell-type-enriched expression of three novel genes in the xenograft single-cell dataset, including Ssc_EHL2chr1g02293 (Endo-enriched), Ssc_EHL2chr17g00009 (PT_VIM-enriched), and Ssc_EHL1chr1g00164 (Immune-enriched). **(c)** Temporal expression patterns of novel genes from bulk RNA-seq of pig-to-human kidney xenograft core biopsies collected at 0, 12, 24, and 48 h post-transplantation, shown as centered log-scaled expression. Symbols denote predicted signal peptides (^*^) or predicted transcription factor domains (^#^), and gene labels are colored by predicted functional categories. **(d)** Genomic organization of the albumin (ALB) locus across haplotype-resolved pig assemblies (hap1 and hap2) and the human reference (T2T-CHM13). Gene models and local sequence context are shown for each assembly. Predicted protein lengths are indicated, and pairwise amino acid sequence identities between the aligned proteins are labeled on the right. **(e)** Genomic organization of the erythropoietin (EPO) locus across pig haplotypes and the human reference. Comparative gene models are shown together with predicted protein lengths, and pairwise amino acid sequence identities between the aligned proteins are labeled on the right. **(f)** A novel secreted protein gene resolved on chrY (hap1) and chrX (hap2) in the T2T assembly. Dashed lines indicate that the locus is located within a PUR. Predicted protein sequences are aligned below to show the high similarity between the two haplotype-specific gene products.

Donor-derived secreted proteins in xenotransplantation can both trigger immune responses and contribute directly to graft function. For example, the kidney-derived hormone EPO regulates erythropoiesis, and although porcine EPO shares substantial sequence similarity with its human counterpart, residual sequence differences (Fig. 4e) may still affect immunogenicity and physiological compatibility in the human host. Because EPO is required for normal red blood cell production, the goal is not simply to eliminate its activity, but rather to preserve or restore a functionally appropriate, human-compatible form. Similarly, after liver transplantation, production of albumin (ALB) is essential because albumin is the major plasma protein responsible for maintaining oncotic pressure and transporting a wide range of endogenous and exogenous molecules. However, porcine ALB also differs substantially from human ALB (Fig. 4d), raising the possibility that cross-species differences in major secreted proteins may influence physiological adaptation after transplantation. In addition, the T2T genome reveals previously unresolved genes encoding organ-enriched secreted proteins that show substantial divergence from their human counterparts (Supplementary Table 14, Fig. 4f). These loci represent additional candidates for future gene-editing strategies aimed at improving graft function and physiological compatibility through pre-transplant humanization of selected secreted proteins.

### Implication of pig T2T Genome in Designing Interspecies Chimerism

A potentially better way to solve human organ shortage is to generate humanized organ in pig through interspecies chimerism^26–31^. Lai and colleagues have managed to generate hiPSC-derived embryonic kidney tissues in pig through gene-edited pig embryos^26^. The availability of high quality T2T genome from both pig and human should allow better design for future attempts. To this end, we integrated organ transcriptomes with cross-species orthology features derived from the haplotype-resolved *Ssc_EHL* assemblies to identify candidate loci relevant to organ-deficient pig engineering and interspecies chimerism. Tissue transcriptomes from heart, kidney, liver, lung, muscle, and spleen have highly concordant expression patterns between haplotypes across all six tissues (Supplementary Fig. 13) and resolved distinct sets of organ-enriched genes on hap1 and hap2. Cross-species orthogroup redundancy analysis indicates that most pig-human orthogroups are close to one-to-one, whereas a subset displays copy-number expansion on either the pig or human side (Supplementary Figs. 14a, b). An ideal target for organ-deficient pig engineering should minimize off-tissue expression while retaining strong pig-human orthology and a simple, well-matched orthogroup structure. To quantitatively prioritize loci potentially relevant to interspecies compatibility, we defined an interspecies chimerism compatibility (ICC) score integrating tissue-specific significance (Si), pig-human genes redundancy (Re), and orthology confidence (Co) (Methods). In each organ, ICC increases with stronger tissue-specific enrichment, higher orthology support and minimal pig-human redundancy imbalance, recovering known lineage regulators among top-ranking examples, including SALL1 and SIX1 for kidney^26^, MYOD, MYF5, and MYF6 for muscle^31^, and HHEX for liver^27^ (Fig. 5a-f). The same organ-resolved ICC patterns are reproduced on hap2, with highly similar distributions of organ-enriched gene sets and ICC landscapes across tissues (Supplementary Fig. 14c-h). For each organ, we report the top 500 ICC-ranked genes and retain loci with one-to-one syntenic or protein-homology support that are shared between hap1 and hap2 as a conservative candidate set for pig-human chimerism studies. (Supplementary Table 15). Genes in these lists include established regulators of the corresponding organs, supporting the biological relevance of the ICC framework. For example, HAND2 in the heart list and PAX5 in the spleen list, consistent with their known developmental roles^32^. We also identified a novel gene (Ssc_EHL2chr5g00233) among the leading candidates in the heart list. This gene was a gene from an unplaced scaffold in *Sscrofa11*.*1* and corresponds to RASD2, further highlighting the value of the T2T genome for refining candidate selection and optimizing gene-editing strategies in pigs.

**Fig. 5.**
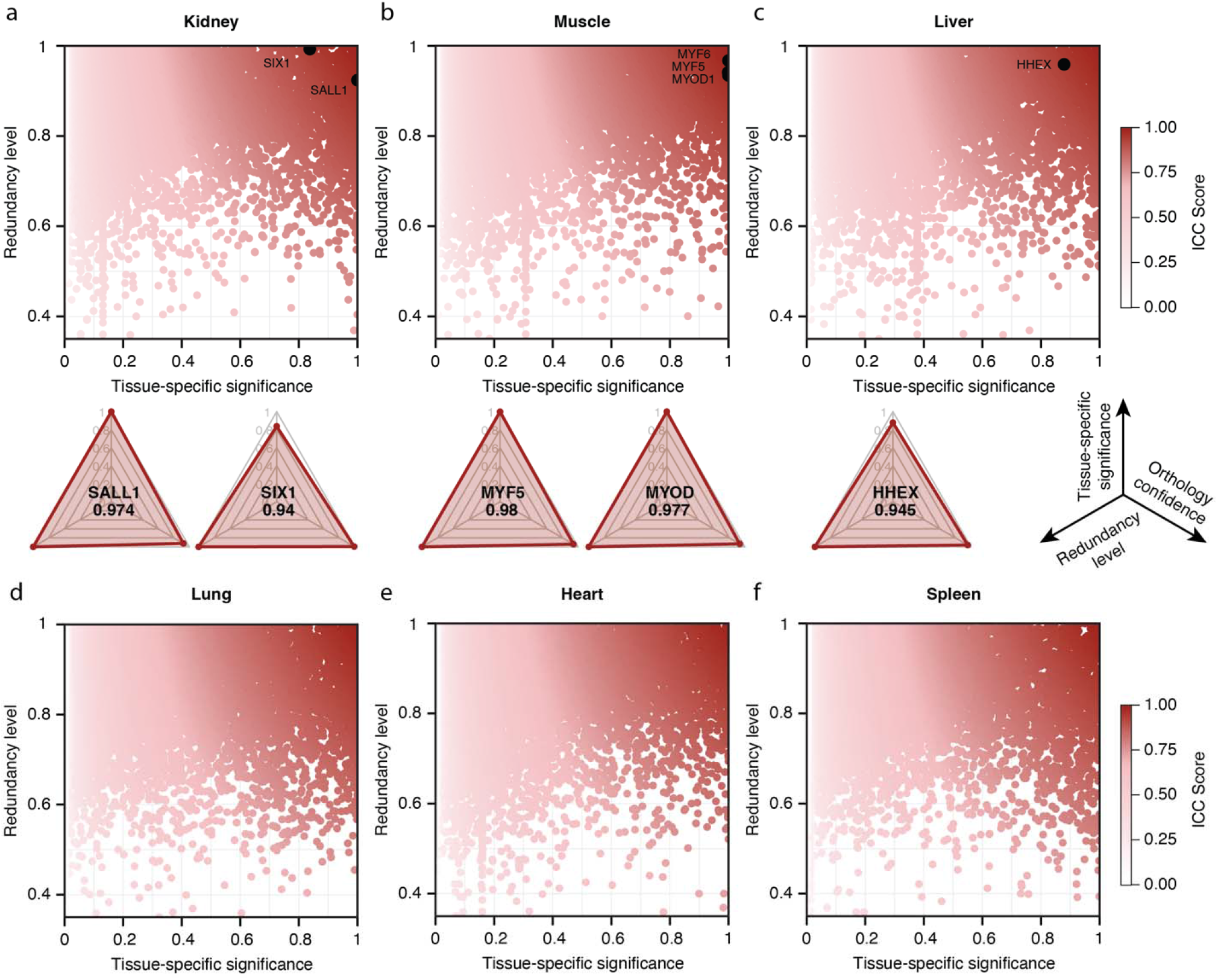
Genomic determinants of interspecies chimerism compatibility. (**a-f)**, ICC score landscape for kidney (a), muscle (b), liver (c), lung (d), heart (e), and spleen (f) of *Ssc_EHL* hap1. Genes are plotted by tissue-specific significance (x-axis) and redundancy (y-axis), with colors indicating ICC scores. Previously reported interspecies chimerism target genes (e.g., *SALL1* and *SIX1* for kidney, *MYOD1, MYF5*, and *MYF6* for muscle, and *HHEX* for liver) are highlighted.

For individual high-ranking genes that may not directly control cell fate, alternative strategies such as Cre-dependent expression of diphtheria toxin subunit A (DTA) driven by gene-specific promoters could be used to achieve targeted ablation and facilitate organ complementation^33^. By integrating tissue specificity, orthology, and redundancy rather than relying solely on expression levels, the ICC framework retains genes with broader expression patterns, avoiding overly restrictive filtering that would exclude functionally redundant or compensatory loci. As a result, candidate genes may overlap across multiple organs, providing a flexible pool for combinatorial design. This enables optimization of gene sets tailored to specific target organs, for example through multi-gene knockout strategies (e.g., SALL1 and SIX1 for kidney) to reduce compensatory effects, or conditional knockout approaches to achieve spatially restricted gene ablation when genes are active in multiple tissues^34^.

Collectively, these analyses provide organ-specific, haplotype-consistent prioritization of candidate loci for interspecies chimerism and organ transplantation by jointly accounting for tissue-restricted expression, pig-human orthogroup redundancy, and protein-level orthology support. This information should be considered prior to the design of any future attempt to generate humanized organ through chimerism.

## Discussion

We report here haplotype-resolved telomere-to-telomere pig genomes for Erhualian (*Ssc_EHL*). The *Ssc_EHL* assemblies resolve both parental haplotypes and enable base-level analyses across centromeres, telomeres, and previously unresolved regions that are typically enriched for repeats and structural variation. These sequences include segmental duplications, high-copy satellite arrays that define functional centromeres, and repeat-associated gene families that are difficult to represent in partially phased references. Notably, the phased T2T assemblies also uncovered previously unmappable PERV loci, many of which located in newly resolved PURs, revealing a more complete and potential breed-level differences of PERV landscape. Together, these components in a phased context provide a more complete view of pig genomic architecture and diversity, and offer a reference framework for regenerative medicine applications, particularly xenotransplantation and interspecies chimerism.

Beyond improving genome completeness, the *Ssc_EHL* assemblies provide a practical framework for linking pig genomic variation to translational applications in these two contexts. By resolving duplicated and repeat-rich loci in a haplotype-resolved manner, it enables more accurate identification of genes and structural features that are often collapsed, misassigned, or incompletely represented in non-phased reference assemblies. In this study, this added resolution recovered previously unresolved gene content, including loci embedded in repetitive sequence space, improved cross-species orthology inference, and identified a set of high confidence pig-specific genes, thereby refining candidate sources of interspecies molecular mismatch. In parallel, applying the haplotype-resolved T2T annotation to public pig-to-human kidney xenograft single-cell transcriptomes enabled finer stratification of injury-associated cell states and uncovered newly resolved, cell-type-enriched genes, improving transcriptomic readouts for evaluating xenograft adaptation and physiological compatibility.

These advances have distinct implications for xenotransplantation and interspecies chimerism. In xenotransplantation, they improve the evaluation of candidate genes potentially relevant to immune incompatibility, xenograft injury and adaptation, and organ physiology. They also enhance interpretation of repetitive and duplicated regions that are difficult to evaluate using earlier references. In interspecies chimerism, complete phasing and more accurate cross-species gene matching between pig and human provide a stronger basis for prioritizing loci relevant to organ development and organ-specific functions in their dosage and haplotype context, which is particularly important for engineering organ-deficient host pigs. Consistent with this, integration of orthology features with organ transcriptomes enabled us to define an Interspecies Chimerism Compatibility score that prioritizes organ-enriched, highly conserved loci with minimal redundancy imbalance, providing a rational framework for nominating engineering targets. Together, these findings show that a haplotype-resolved T2T pig genome is not merely a more complete reference, but an evidence-based genomic foundation for systematic cross-species comparison, improved interpretation of repetitive loci, and target prioritization in xenotransplantation and interspecies chimerism.

## Methods

### Sample collection and sequencing

Fresh whole-blood samples were collected from a healthy Erhualian sow and boar at Changshu Agriculture, Industry and Commerce Company, Limited. Tissue samples (muscle, heart, lung, spleen, liver, kidney) were collected from a 73-day-old castrated male pig from the same facility. Blood samples were collected into EDTA-containing blood collection tubes, and immediately frozen on dry ice. After slaughter, tissue samples were rapidly frozen in liquid nitrogen and stored at −80⍰°C until further analysis. DNA and RNA extraction, library construction, and sequencing were performed at GrandOmics, including PacBio HiFi WGS and Iso-Seq on the PacBio Revio, Oxford Nanopore ultra-long WGS and Pore-C on the PromethION, and MGI WGS and RNA-seq on the DNBSEQ-T7RS.

### Genome Assembly and Haplotype Resolution

#### Assembly methods for paternal and maternal haplotypes

Two kmer databases for parents were built with yak (version 0.1-r66-dirty, parameters: -b 37 -k 21) from 30X whole-genome sequencing data collected from the parental and maternal pigs, respectively, and filtered with fastp (v0.20.0) using default parameters. Two primary genome assemblies containing parental/maternal source contigs were obtained using hifiasm (version 0.25.0, parameters: --ont -trio-dual) from ONT reads longer than 50kb and a parental/maternal kmer database. We aligned the primary assembled contigs to the pig reference genome (Sscrofa11.1) and found that the assemblies of chromosomes X and Y were already gapless, telomere-containing, and of proper size. We separately collected the ONT reads mapped to parental or maternal genomes and performed additional assembly using HiFiasm with the parameters −3 −4, resulting in primary assemblies of the parental and maternal genomes.

The quality-passed Pore-C sequencing data were independently aligned to two primary assemblies with minimap2(version v2.26, parameters: -x map-ont -B 3 -0 2 -E 5 -k13 -a), and paired BAM files were generated from the PAF files, the mapping results were further filtered according to following criteria: mapping quality ≥ 1, alignment_length >= 90, and percent_identity >= 150. The contigs from two genomes were clustered respectively with haphic (v1.0.5) applying the Markov Cluster Algorithm (MCL algorithm) by default parameters. The 3D-DNA iterative scaffolding algorithm was subsequently used to rapidly sort and orient the contigs. The contigs were mapped to chromosomes, resulting in chromosome-level genome sequences.

To fill the gaps between contigs mounted through Pore-C, we aligned Nanopore Ultra Long reads (Super Basecalling modeler) to the mounted genomes, then extracted the unmapped reads and reads uniquely mapped to the dual sides of a gap, and performed local iterative assembly using nextdenovo to find the best gap-filling path, the assembled sequences matching the end of gaps were aligned back to the corresponding gaps and replace the gap regions.

After filling the gaps, the two genome assemblies were polished to the final version using Nextpolish2, based on 21k-mer and 31k-mer libraries constructed from quality-controlled whole-genome sequencing data, and a BAM file generated by aligning HiFi reads to the gap-free genomes using minimap2 (parameter: -map-hifi).

To identify the telomere of each chromosome, we apply the seqkit locate software to search for the telomere repeat unit AACCCT. For the chromosomes lacking the telomere sequence at the end, a local assembly was performed using HiFiasm from three types of HiFi reads: (1) reads unmapped to the genome, (2) reads containing telomeric sequence (AACCCT), and (3) reads mapped to chromosomes not mapped with a telomeric region within 1Mb. The newly assembled contigs were connected to the telomere-lacking chromosomes by identifying overlaps between the contigs and the chromosome ends using minimap2 to obtain the complete chromosomes.

#### Assessment of assembly quality and completeness

A 21-mer database was generated from HiFi sequencing data using Merfin. The QV (Quality Value) and error rate statistics for each chromosome and the whole gap-free genome were calculated using Merqury based on the k-mer evaluation method to validate the assembly quality. The completeness of assembly was evaluated through predicting the gene content of the existing sequences in the assembled genome from the OrthoDB database and calculating the BUSCO (Benchmarking Universal Single-Copy Orthologs) score.

### Centromere Analysis

#### Identification and characterization of centromeric and telomeric regions

The candidate high-repeat regions of the gap-free genome were identified in parallel by the TRF (v4.09.1) with default parameters and the RepeatMasker (v1.331) with parameters nolow -no_is -gff -norna -engine abblast -lib lib, and the corresponding sequences were extracted using samtools (v1.3). 27-mer frequencies within these regions were counted using KMC (v3.2.4) with -k27 -m512 -t60 -ci100 -cs1000000, and k-mer tables were exported using kmc_dump. Candidate satellite monomers were assembled from k-mer tables using SRF (https://github.com/lh3/srf). Each satellite candidate was mapped to the gap-free assemblies using BLASTN (v2.14.1+), and hits were sorted by genomic coordinate. For each candidate, we summarized chromosome-scale hit distributions and inter-hit distances; candidates showing long runs of regularly spaced hits were considered tandem arrays representative of centromeric satellites.

To thoroughly investigate the sequence organization of the centromere of each chromosome, we applied the TRASH pipeline (https://github.com/vlothec/TRASH_2) to identify all satellite monomers with --maxdiv 10 --par 60. The 335 ± 10 bp satellite monomers assembled by SRF were used as TRASH templates. Candidate centromere intervals were defined as the longest loci on each chromosome containing continuously distributed short repeats, with a maximum gap less than 10 kb between adjacent repeat elements. To characterize satellite subfamilies and their chromosomal distributions, all TRASH-annotated monomers matching the 335 ± 10 bp templates were extracted (229,672 monomers). Monomers occurring more than 5 times (87,590 instances; 6,208 unique sequences) were aligned with MUSCLE (v3.8.1551), and a phylogenetic tree was inferred using FastTree (v2.2) with -nt -gtr. Based on the phylogenetic tree, 12 satellite subfamilies were manually delineated; subfamily-assigned sequences were extracted from the MUSCLE alignment, and sequence logos were generated with WebLogo (v3.7.12) after filtering low-support positions (<5% frequency). Remaining unclassified monomers were assigned by BLASTN to the closest subfamily representative and labeled accordingly.

#### CENP-A CUT&RUN sequencing and analysis

CUT&RUN library preparation was performed by Hyperactive pG-MNase CUT&RUN Assay Kit for Illumina (Vazyme HD102) according to a published protocol^35^. Ear fibroblasts were cultured in DMEM (Gibco C11995500BT) with 15% FBS (Gibco 10099141C), supplemented with 1% nonessential amino acids (Gibco 10565018) and 1% GlutaMAX (Gibco 35050061). Cells were confirmed mycoplasma-free detected by Mycolor One-Step Mycoplasma Detector (Vazyme Biotech Co.,Ltd D201). Briefly, 100,000 cells were immobilized on Concanavalin A-coated magnetic beads and permeabilized with digitonin. Cells were incubated with a CENP-A primary antibody (CST 2168) at 1:50 against the target protein at 4°C overnight, followed by binding of pG-MNase fusion enzyme. DNA cleavage was carried out on ice for 60 min upon Ca^2+^ activation. The reaction was stopped by adding Stop Buffer with spike-in DNA, and released DNA fragments were purified using spin columns. Libraries were constructed through end repair, adapter ligation, and PCR amplification for 14 rounds, then validated for size distribution and sequenced on the Illumina NovaSeq X plus PE150 platform.

CUT&RUN reads and a published pig CENP-A ChIP-seq dataset (SRA: SRR30630686) were adapter-trimmed using Trim Galore (v0.6.4). Trimmed reads were aligned to the gap-free assemblies using bowtie2 (v2.5.4) with --very-fast --no-mixed --no-discordant -k 10. Alignments were filtered to remove mitochondrial reads and PCR duplicates using Picard. CENP-A signal tracks were generated from filtered BAM files using deepTools bamCoverage (v3.3.2) with 50 bp bins and RPKM normalization, and exported as bigWig files.

### Genomic Structural and Variational Analysis

#### PUR Detection

PURs were detected as previously described by Liu *et al*.^*7*^. Each reference assembly (*Sscrofa11*.*1, Ssc_WZS, Ssc_JH, Ssc_RC, Ssc_MIN*) was aligned to the *Ssc_EHL* assemblies using winnowmap2^36^ (v2.0.3) with parameters -ax asm20 -H -MD -t 16. SAM alignments were converted to PAF format using paftools^37^. Unaligned (or low-confidence) intervals on the *Ssc_EHL* assemblies were extracted from the PAF alignments using bedtools (v2.31.0), defining PURs as regions lacking confident alignments. For visualization, PUR intervals larger than 50 kb were retained and plotted along chromosomes using karyoploteR^38^ (v1.24.0).

#### Detection of structural variations (SVs) and copy number variations (CNVs)

Structural variants (SVs) and copy-number variations (CNVs) were identified by whole-genome alignment with minimap2^39^ (v2.29-r1283) followed by SyRI^40^ (v1.7.1). *Sscrofa11*.*1, Ssc_WZS, Ssc_JH, Ssc_RC*, and *Ssc_MIN* assemblies were used as query genomes against *Ssc_EHL* assemblies. Counts and size distributions of SVs/CNVs were summarized and visualized in GraphPad Prism (v10.6.1). Synteny and rearrangement plots were generated using plotsr (v1.1.1).

### Repeat Annotation

Repeat annotation of the gap-free genome assembly was performed using RepeatMasker (v4.1.7, http://www.repeatmasker.org) with a customized repeat database to improve the sensitivity and accuracy of transposable element (TE) detection. The default repeat library provided by RepeatMasker was replaced with a specific Dfam database version (Dfam_3.8), which contains abundant, high-quality multiple sequence alignments and consensus sequences for known repetitive elements in pig.

### Gene Annotation and Functional Genomic Analysis

#### Initial gene annotation and refinement

Gene models were generated on a repeat-masked genome by integrating (i) homology-based prediction with GeMoMa using peptide alignments from related species, (ii) RNA-seq supported models by mapping reads with STAR followed by transcript assembly with StringTie and ORF/UTR inference with PASA, and (iii) ab initio prediction with AUGUSTUS trained on PASA-derived transcripts. For downstream analysis, *Sscrofa11*.*1* gene models were projected onto each T2T haplotype using Liftoff (v1.6.3) and miniprot (v0.18). Liftoff was run with - exclude_partial and -polish to remove partial or low-identity mappings and to polish exon structures by realignment. Miniprot-derived models were reconciled locus-by-locus with the draft T2T annotation and Liftoff projections. For loci supported by multiple sources, models with CDS coverage ≥70% were considered concordant, and the best-supported model (draft T2T, Liftoff, or miniprot) was retained based on CDS coverage and exon integrity; the retained source label was recorded in the final GTF. Miniprot-supported loci that did not overlap any draft T2T/Liftoff gene model were retained as “Unanchored” loci. Draft T2T loci lacking support from both Liftoff and miniprot were retained as “Novel” only if they met structural criteria (≥3 exons and CDS ≥200 bp) and showed evidence of expression in at least one of the six organs or functional annotation support, and were labeled accordingly in the final GTF.

#### Characteristics of novel genes

Amino acid sequences were obtained from CDS regions of novel gene models and searched against the *Sscrofa11*.*1* protein set and the reviewed UniProt/Swiss-Prot database using BLASTP. Transcription factor domains were predicted using AnimalTFDB. SignalP 6.0 was used to predict signal peptides, with haplotype1 and haplotype2 protein sets analyzed separately using the eukaryotic model. It reports the predicted probability and the signal-peptidase cleavage site. Functional annotations for novel genes were inferred using SPROF-GO^41^, which predicts Gene Ontology (GO) predictions across Molecular Function (MF), Cellular Component (CC), and Biological Process (BP) from the protein sequences. For each ontology, we retained the top 10 predicted terms per gene and applied a confidence threshold to removed low-confidence terms while ensuring at least one prediction. Detailed characteristics are provided in Supplementary Table 10.

### Gene Expression Reanalysis during Early Embryogenesis

#### Stage-specific gene expression

Public bulk RNA-seq (CRA006174) and scRNA-seq (CRA003960; GSE112380) datasets spanning porcine pre-implantation development were collected. Differential expression in bulk RNA-seq was assessed with DESeq2 (v1.42.0). For scRNA-seq, stage-enriched genes were identified using Seurat (v5.3.0) with FindMarkers (avg_log2FC > 1, adjusted P < 0.05). Genes enriched at the 2-cell, 4-cell, and 8-cell stages were defined as early-stage specific genes (ESGs). ICM-specific genes (ISGs) were defined as genes enriched in ICM relative to TE at E5, and epiblast-specific genes (EpSGs) were defined as genes enriched in epiblast relative to hypoblast at E7.

#### Bulk RNA-seq analysis

Bulk RNA-seq reads were trimmed with Trim Galore (v0.6.4) and aligned to the *Ssc_EHL* genome using HISAT2 (v2.2.1). Gene-level expression was quantified from aligned BAM files using GFOLD (v1.1.4) with *Ssc_EHL* annotations, and gene-wise values were used for downstream comparisons.

#### Single-cell RNA-seq data analysis

Raw reads from scRNA-seq were demultiplexed using 8-bp cell barcodes in Read 2, allowing up to two mismatches. 8-bp UMIs in Read 2 were appended to the corresponding Read 1 headers using UMI-tools (v1.1.6). Read 1 was trimmed to remove template-switch oligo (TSO) sequences, poly(A) tails, and low-quality bases using Cutadapt (v3.7). Processed reads were grouped by cell barcode and aligned to the final reannotated genome with HISAT2 (v2.2.1). Aligned reads were assigned to annotated genes, and gene-level values were quantified using GFOLD (v1.1.4).

### Cross-species Synteny Analysis

Protein sequences were obtained for human (T2T-CHM13v2.0, GCA_009914755.4), mouse (T2T_mhaESC_v1.0, GCA_050437135.1), and *Ssc_EHL* assemblies. For each gene, the longest protein-coding isoform was retained. Orthology and synteny blocks were inferred using the jcvi toolkit^42^ (v1.5.9) with default parameters, integrating protein homology with gene order information.

### Pig-Specific Gene Detection

Pig-specific genes were identified using a stepwise filtering pipeline. First, orthogroups were inferred with OrthoFinder (default settings) using human proteins and *Ssc_EHL* proteins. Pig genes not assigned to any human-containing orthogroup were retained as initial candidates. Candidates were refined by reciprocal BLASTp searches against the human proteome using default parameters. Genes were retained as BLASTP-negative if they showed weak similarity in both directions (E-value > 1e-3 or bit score < 70). Finally, human-pig synteny was assessed using jcvi. Ortholog candidates were generated with jcvi.compara.catalog ortholog, and synteny anchors were filtered using jcvi.compara.synteny screen --simple. Refined candidates not falling within any conserved synteny block were designated as pig-specific genes.

### Interspecies Chimerism Compatibility Score

To quantitatively prioritize loci for interspecies chimerism, we integrated three metrics capturing tissue specificity, redundancy balance of pig-human, and orthology confidence. (1) *Tissue-specific significance* (*Si*), which quantifies the expression specificity of a gene in a given tissue relative to others, derived from GFOLD values across tissues. For gene *j*, Si is defined as the mean value of {0, *v*_*j*_}_*max*_ over all tissues, where *v*_*j*_ denotes the gfold value of gene *j*. (2) *Redundancy level of genes* (Re), derived from orthogroups to quantify cross-species copy-number imbalance. For each pig-human orthogroup mapping, *n*_*pig*_ and *n*_*human*_ denote the numbers of pig and human genes in this orthogroup. (3) *Orthology Confidence* (*Co*), assessed using OrthoFinder to measure similarity between homologous proteins in the human and pig genomes. The Co score of gene *j* is defined as the maximum of the pig-to-human and human-to-pig similarity scores. Given the distinct scales and distributions of these metrics and no evidence based weights were available, the interspecies chimerism compatibility (*ICC*) score was defined as the geometric mean of the scaled components.

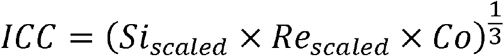

The *Si*_*scaled*_ is derived from *tanh*(*c*_l_ × *Si*), where *c*_1_ is an adjustment parameter to set the scaled level. *Re*_*scaled*_ is calculated by exp (*c*_2_ × |*n*_*pig*_ − *n*_*human*_ |(*n*_*pig*_ − 1)), which emphasizes that suitable candidate genes should be preferentially selected from those with a relatively small copy number of porcine homologs and a minimal difference in copy number between human and pig. The maximum is 1 when *n*_*pig*_ = *n*_*human*_ = 1. The parameter *c*_2_ adjusts the penalty degree for copy number differences. Since *CO* naturally falls within the interval [0,1], no additional scaling is applied. Smaller values of *c*_1_ and *c*_2_ lead to a more stringent scoring scheme. Parameters were set to *c*_1_ = *c*_2_ = 0.5. Genes with high *ICC* values were prioritized as candidate genes and *ICC* scores were computed independently for hap1 and hap2. For each organ, we reported the top 500 *ICC*-ranked genes and retained the one-to-one syntenic blocks shared between hap1 and hap2 as a conservative set of candidate loci for pig-human chimerism studies

## Data availability

The raw sequencing data have been deposited in the NCBI database under the accession number PRJNA1416575.

## Acknowledgements

This work was supported by the Noncommunicable Chronic Diseases - National Science and Technology Major Project (2025ZD0552201), the National Key Research and Development Program of China (2025YFA1804200), the Zhejiang Provincial Natural Science Foundation of China (LZ25C060003), the Yangtze River Delta Sci-Tech Innovation Community Joint Research Project (2022CSJGG1000), the “Pioneer” and “Leading Goose” R&D Program of Zhejiang (2025C01115). We thank the High-Performance Computing Center of Westlake University for computation support. We also appreciate the help from all the members of Cell Fate Control Lab.

## Author contributions

C.C.Z. conceived the study, designed the overall analytical strategy, performed data analyses, secured funding, and drafted the manuscript. Z.T.Z., Z.J.L., J.J.L., W.S., and Y.X.Y. performed computational analyses, generated visualizations, and revised the manuscript. Y.H.W., M.X.S., T.C.K., and B.W.^3^ contributed to data analyses. B.B.S., X.M.W., J.Z.X., and C.J.X., contributed to sample collection and processing. Y.F., J.M., Y.Q., and J.Q.K. contributed to project design and discussions. H.N.W. coordinated and oversaw sample acquisition. B.W.^3,4^ coordinated resources and external collaborations. D.Q.P. supervised and conceived the whole study, revised the manuscript, and secured funding. All authors reviewed and approved the final manuscript.

## Declaration of generative AI and AI-assisted technologies in the manuscript preparation process

During the preparation of this work the authors used ChatGPT for language editing and polishing. After using this tool, the authors reviewed and edited the content as needed and take full responsibility for the content of the published article.

## Supplementary Figures

**Supplementary Figure 1.**
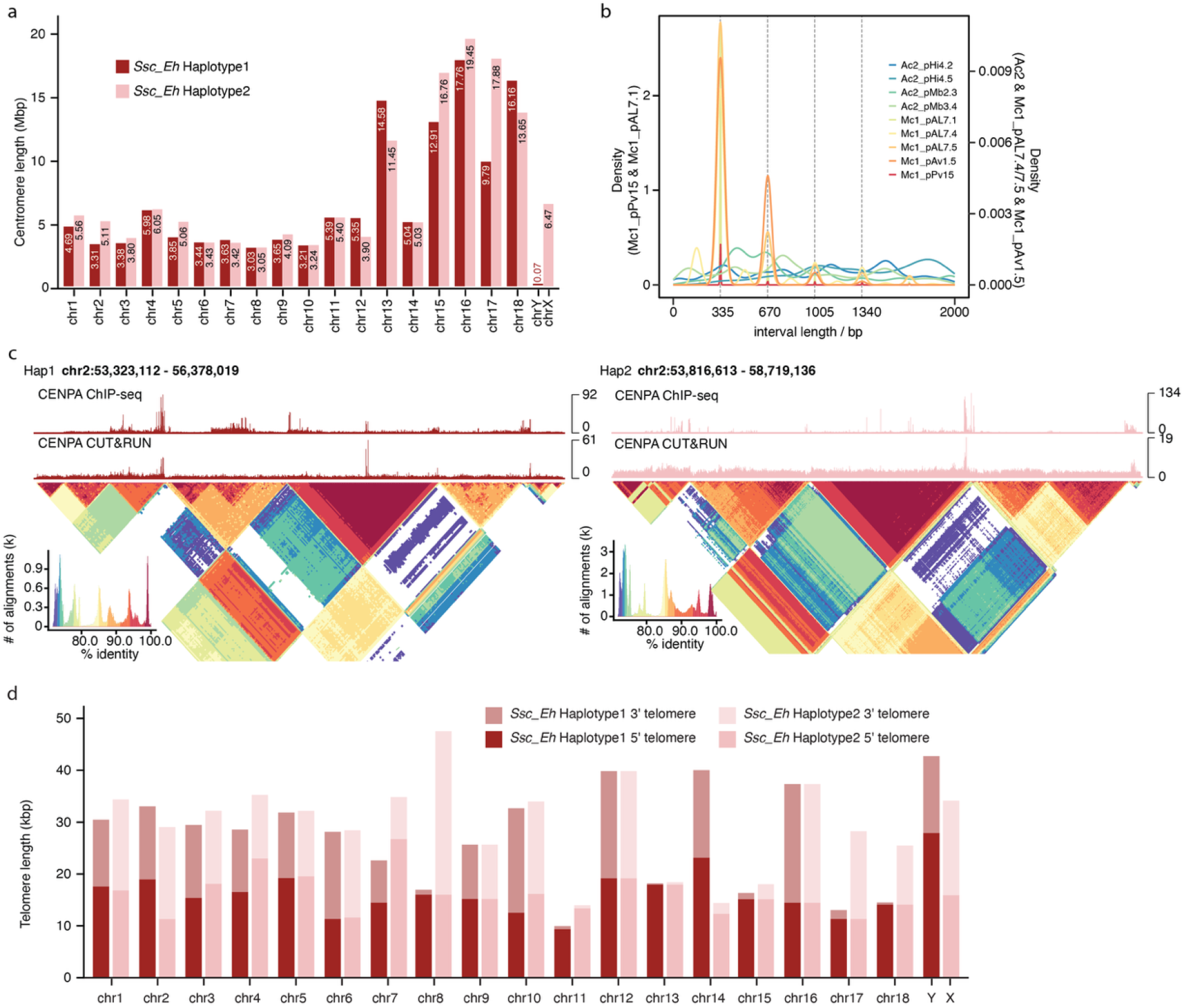
Centromere and telomere identification in the haplotype-resolved T2T Erhualian pig genome.. **(a)** Length of annotated centromeric regions across all chromosomes for both *Ssc_EHL* haplotypes. **(b)** Length distributions of major tandem-repeat candidates corresponding to previously reported centromeric repeat segments (Mcs and Acs), highlighting the dominant ~335 bp satellite monomer. **(c)** Representative centromeric locus on chromosome 2 in hap1 and hap2, showing enrichment of CENP-A ChIP-seq and CENP-A CUT&RUN signals over the reconstructed satellite array. Self-alignment heatmaps and sequence identity histograms summarize the internal homology and higher-order organization of the array. (d) Estimated telomeric tract lengths at the 5′ and 3′ ends of each chromosome for both haplotypes, confirming telomere representation at both ends of all assembled chromosomes.

**Supplementary Figure 2.**
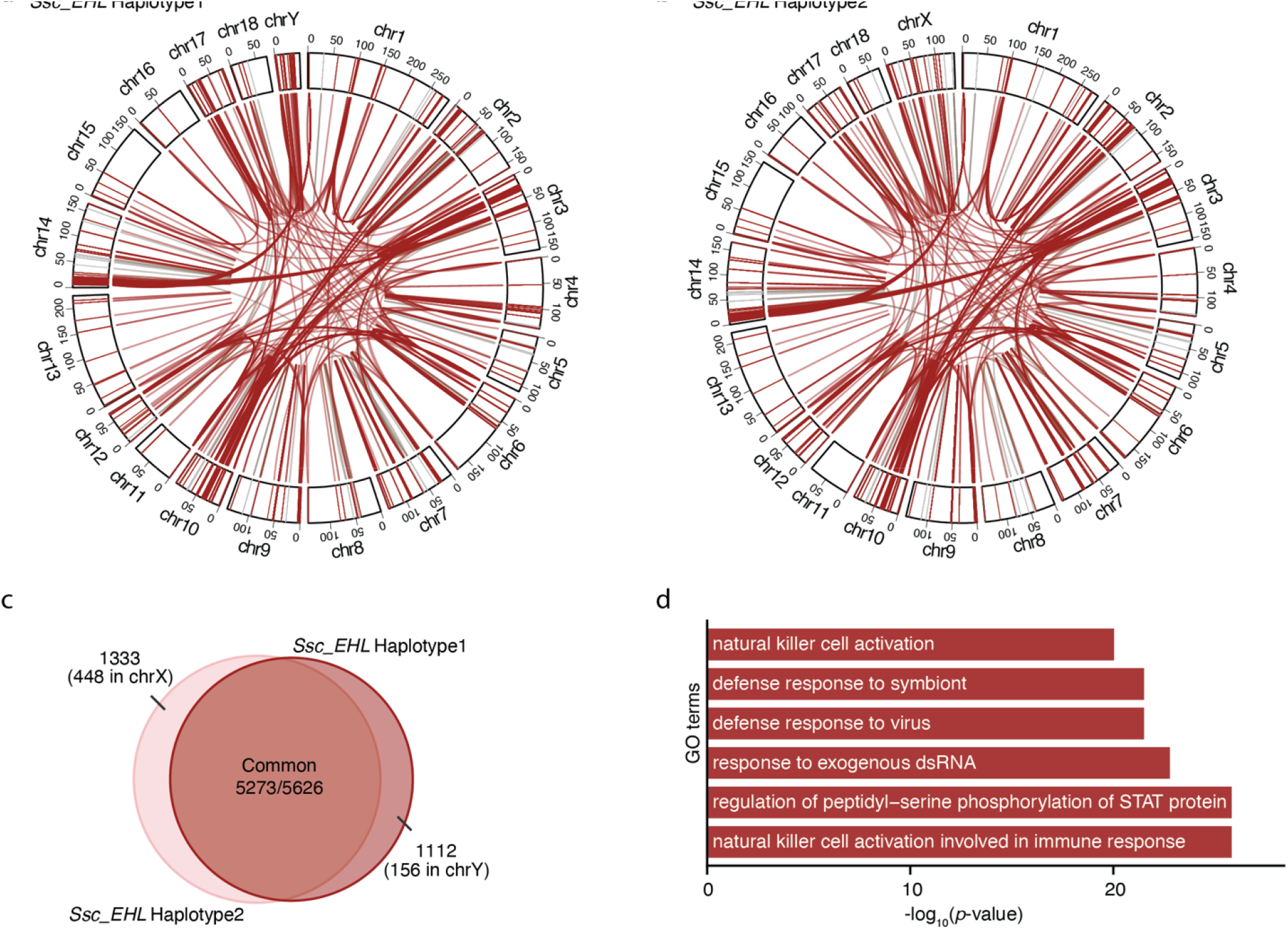
Segmental duplications features in the *Ssc_EHL* genome. (**a-b)** Circos plots highlighting the previously unresolved genome-wide SDs (red line) for hap1 (a) and hap2 (b). **(c)** Haplotype overlap of SDs. Intersection indicates shared SDs and non-overlapping segments are haplotype-specific. **(d)** Gene Ontology enrichment analysis for genes within shared SDs.

**Supplementary Figure 3.**
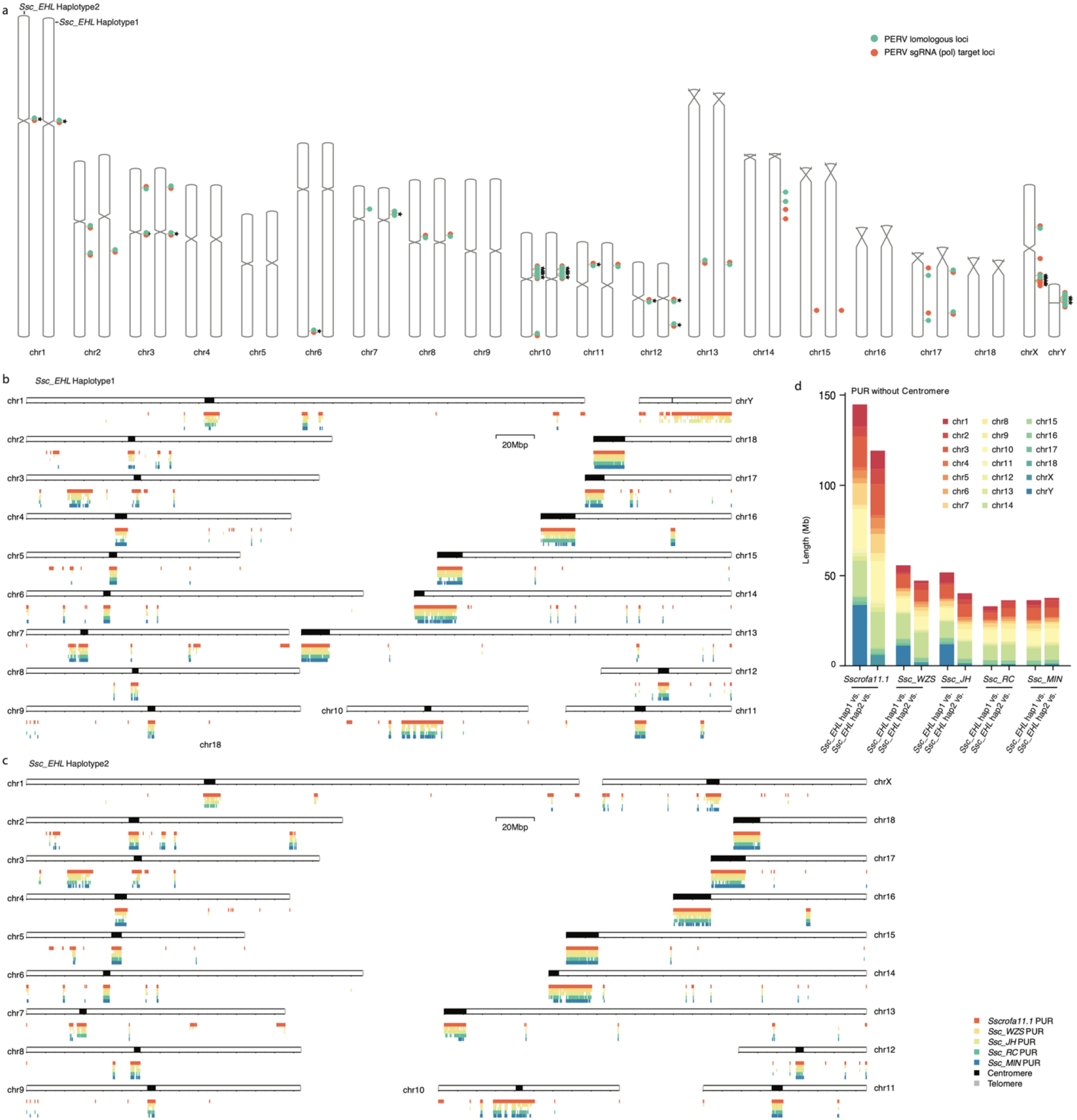
Genome-wide distribution of PURs and non-centromeric PUR burden across pig assemblies. **(a)** Chromosomal ideograms show the positions of PERV homologous loci (green) and PERV sgRNA target loci (orange) across both haplotypes. Stars indicate loci located within pig-unresolved regions (PURs). **(b-c)** Chromosome ideograms of Ssc_EHL hap1 (**b**) and hap2 (**c**) showing the genomic positions of PURs identified in comparisons to Sscrofa11.1 and near-T2T pig assemblies (Ssc_WZS, Ssc_JH, Ssc_RC, and Ssc_MIN). Coloured segments mark PUR intervals for each comparison; centromeres and telomeres are indicated on the ideograms. Scale bar, 20 Mb. **(d)** PUR length after excluding centromeric regions (“PUR without Centromere”) in each comparison using Ssc_EHL hap1 or hap2 as reference. Bars are stacked by chromosome to show genome-wide distribution of non-centromeric PURs across assemblies.

**Supplementary Figure 4.**
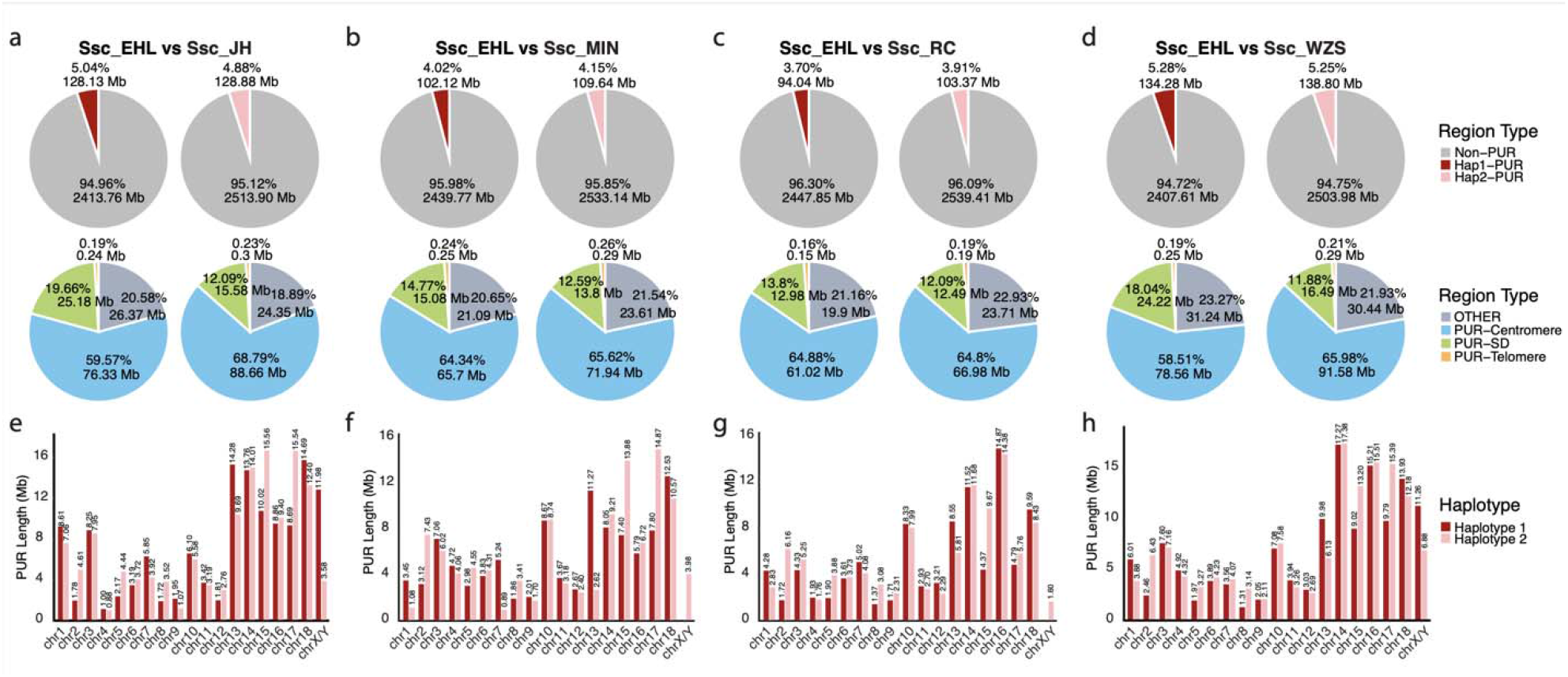
Pairwise PUR composition and chromosome-level PUR profiles in inter-breed comparisons. **(a-d)**, Pairwise comparisons between Ssc_EHL and previously published pig assemblies: Ssc_JH (**a**), Ssc_MIN (**b**), Ssc_RC (**c**), and Ssc_WZS (**d**). For each target assembly haplotype (left and right pies per panel), the upper pie charts show the fraction o aligned sequence (non-PUR) versus PURs attributable to Ssc_EHL hap1 or hap2. The lower pie charts partition the total PURs into centromeric, SD-associated, telomeric, and other components. **(e-h)** Chromosome-wise PUR lengths (Mb) for Ssc_EHL hap1 and hap2 in comparisons to Ssc_JH (**e**), Ssc_MIN (**f**), Ssc_RC (**g**), and Ssc_WZS (**h**).

**Supplementary Figure 5.**
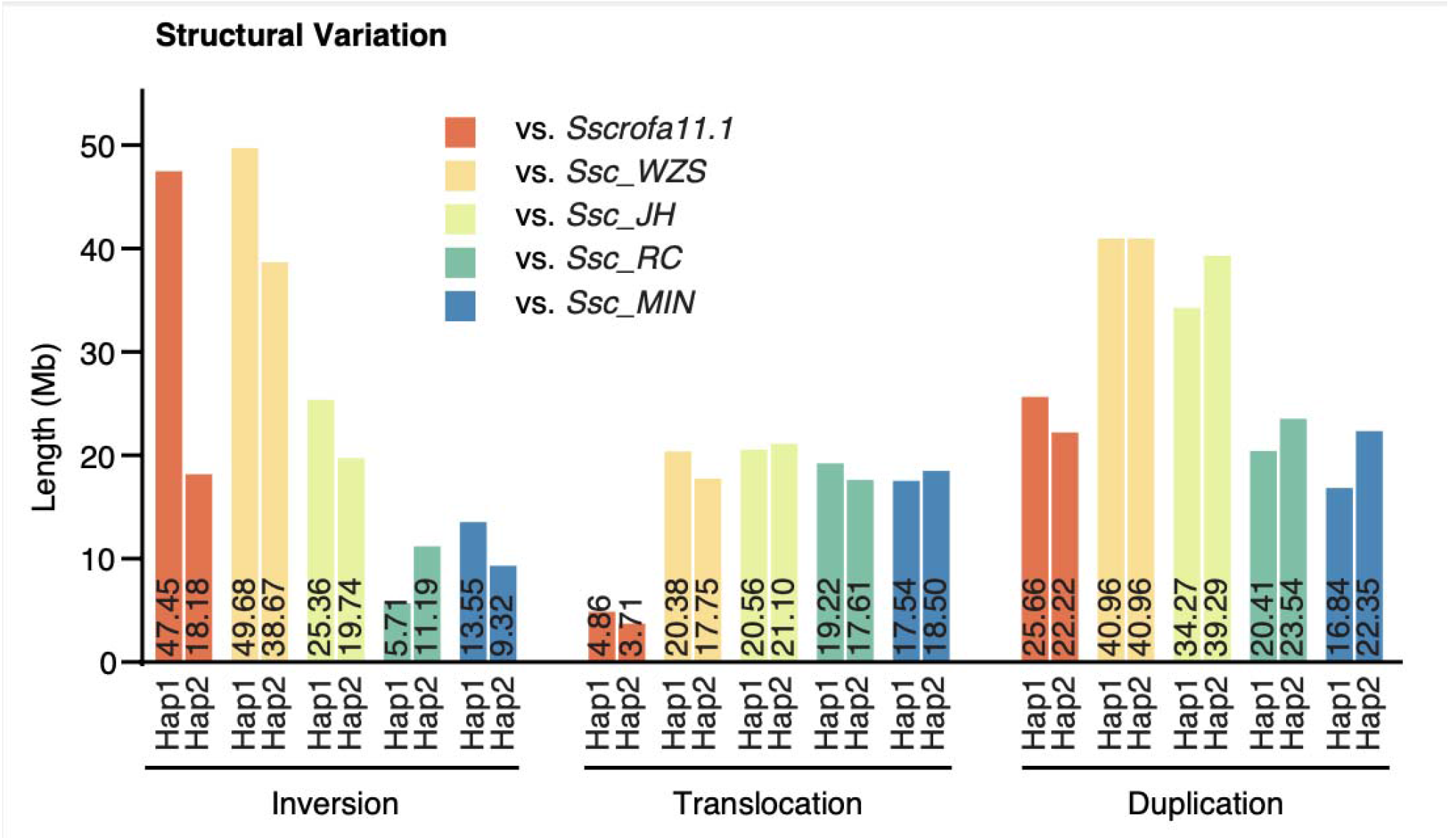
Structural variation burden across pig assemblies using Ssc_EHL haplotypes as references. Total lengths (Mb) of inversions, translocations, and duplications identified when comparing Ssc_EHL hap1 and hap2 to Sscrofa11.1 and previously published pig assemblies (Ssc_WZS, Ssc_JH, Ssc_RC, and Ssc_MIN). Bars are grouped by SV class, with colours indicating the comparison assembly and values annotated on bars.

**Supplementary Figure 6.**
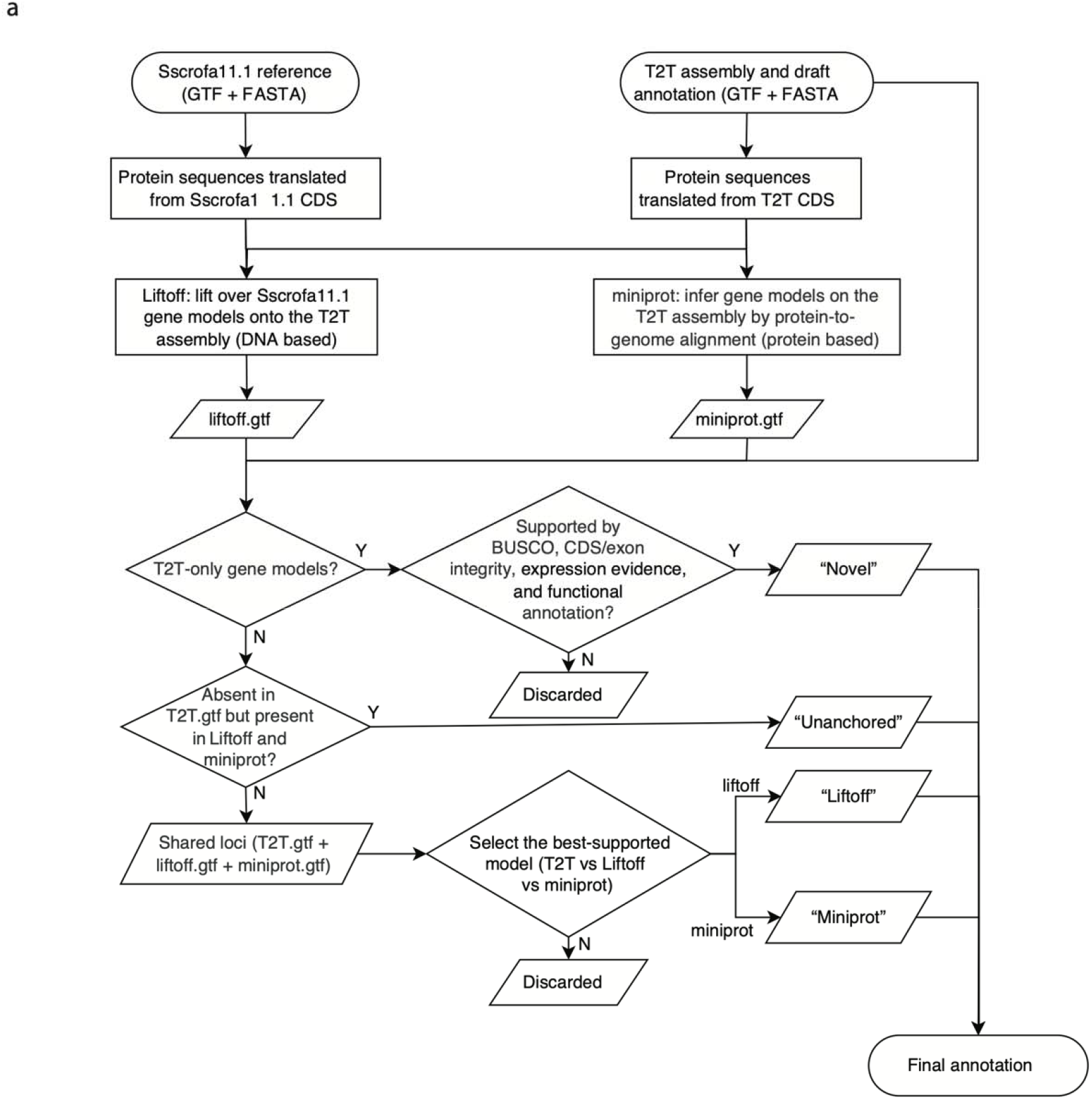
Workflow for reconciling gene models across annotation sources. **(a)** Schematic of locus-by-locus model integration using the Sscrofa11.1 reference (GTF/FASTA) and the T2T assembly with draft annotation. Liftoff projects reference gene models onto the T2T assembly, and Miniprot infers protein-to-genome alignments. At shared loci, the best-supported model is retained; well-supported T2T-only loci are classified as Novel; loci supported by Liftoff/Miniprot but absent from the draft set are retained as Unanchored; unsupported or conflicting models are discarded.

**Supplementary Figure 7.**
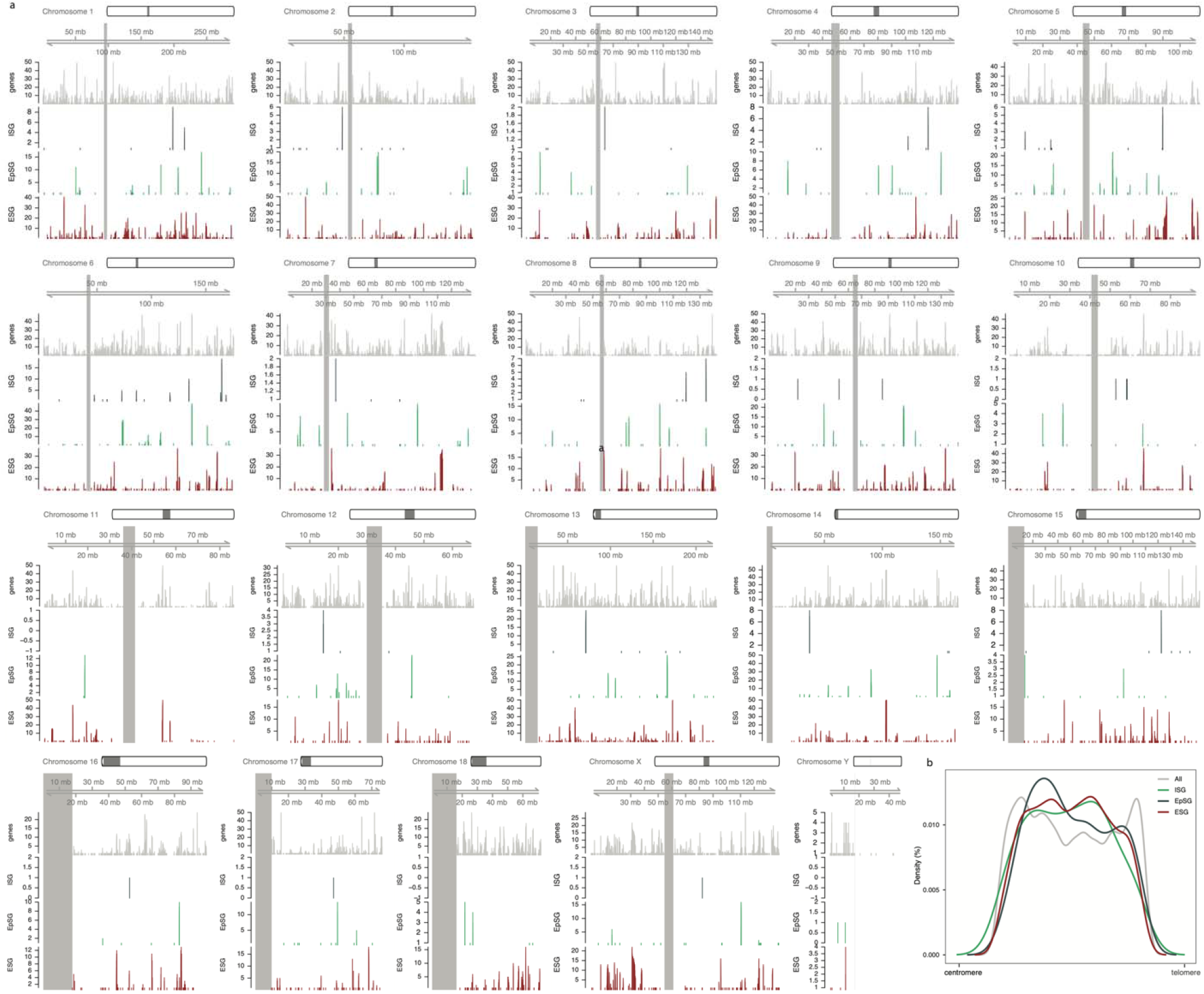
Genome-wide localization of stage-specific gene sets in the T2T assembly. **(a)** Chromosome-level distributions of ESGs, ISGs, and EpSGs along Ssc_EHL chromosomes, shown together with overall gene density; shaded regions indicate centromeric intervals. **(b)** Density of gene positions along normalized chromosomal coordinates from centromere to telomere for all genes and each stage-specific set, highlighting positional enrichment patterns.

**Supplementary Figure 8.**
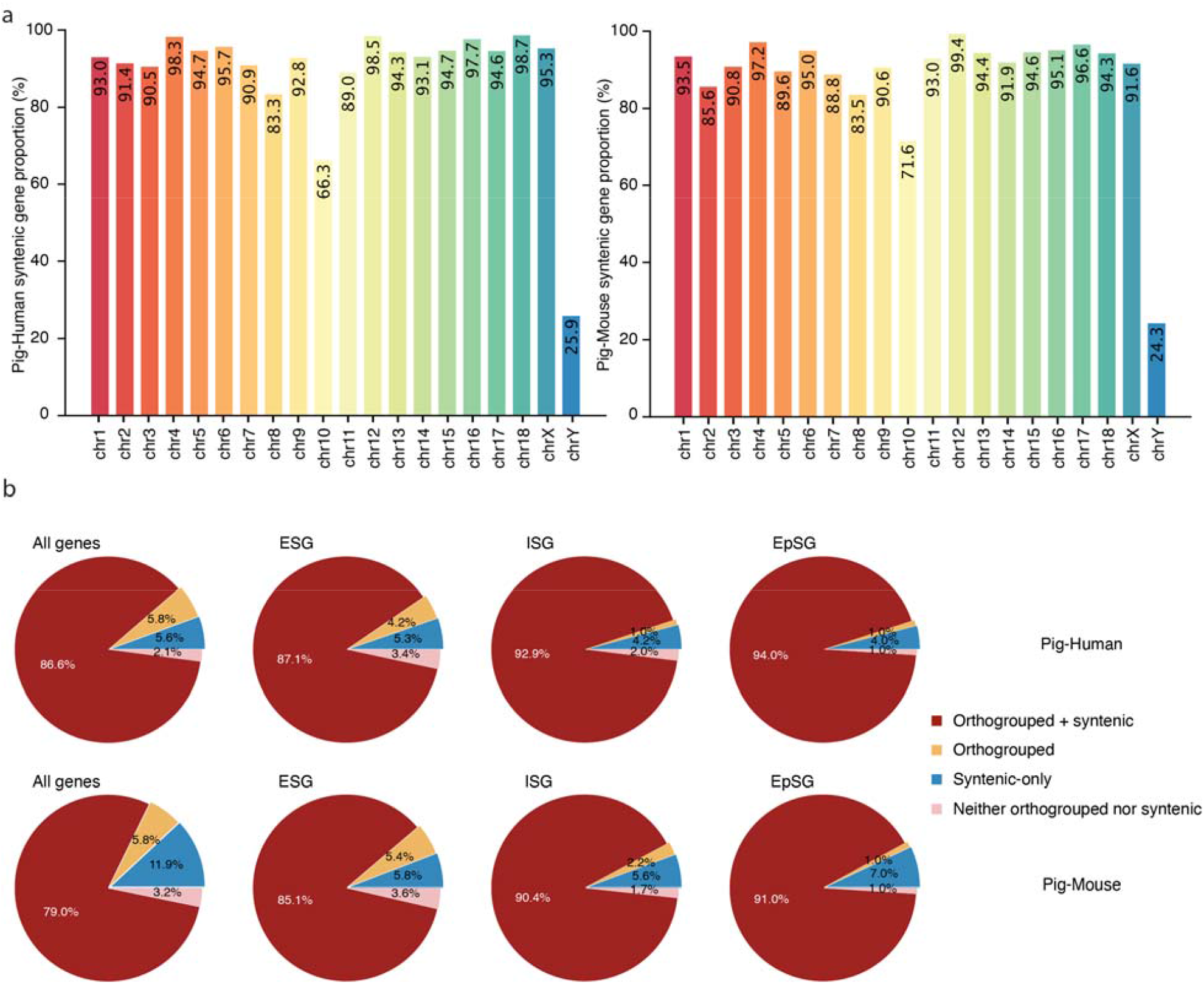
Synteny coverage and combined orthology-synteny support across chromosomes and gene sets. **(a)** Per-chromosome proportion of pig genes assigned to conserved syntenic regions in pig-human and pig-mouse comparisons. **(b)** Partition of pig genes into four categories integrating orthogroup assignment and synteny support (orthogrouped and syntenic, orthogrouped-only, syntenic-only, neither) for all genes and for ESGs, ISGs, and EpSGs, shown separately for pig-human and pig-mouse.

**Supplementary Figure 9.**
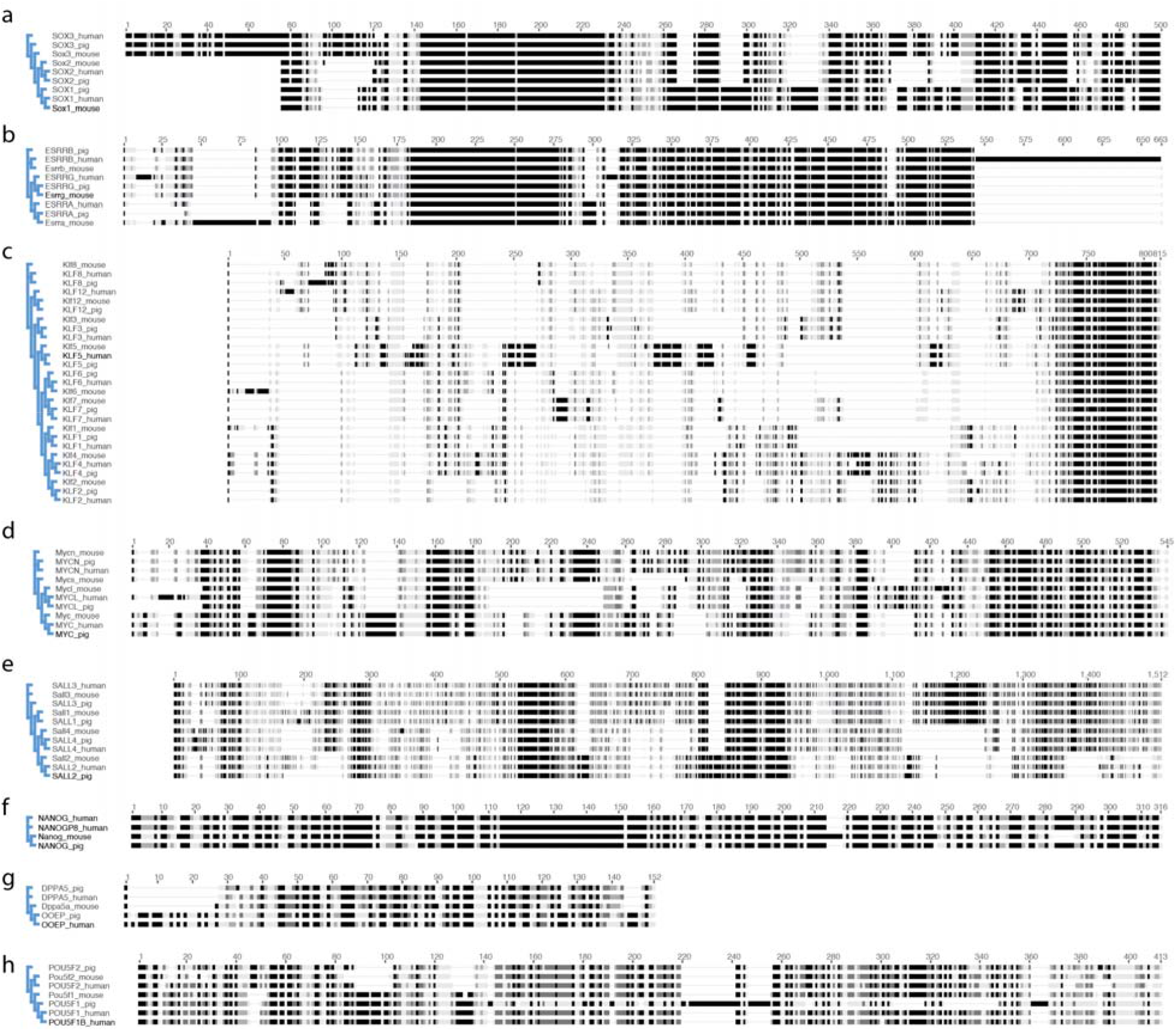
Protein-level alignments of representative pluripotency genes (families) with divergent conservation patterns. Multiple-sequence alignment summaries for selected gene families, highlighting species-specific expansion and divergence. Panels show alignment profiles for **(a)** SOX family members, **(b)** ESRRB/ESRR family, **(c)** KLF family, **(d)** MYC family, **(e)** SALL family, **(f)** NANOG, **(g)** DPPA5, and **(h)** POU5F1/POU5F1-related proteins across pig, human, and mouse. The phylogenetic trees generated by neighbor-joining method are displayed on the left and horizontal tracks summarize aligned blocks and gaps across each protein.

**Supplementary Figure 10.**
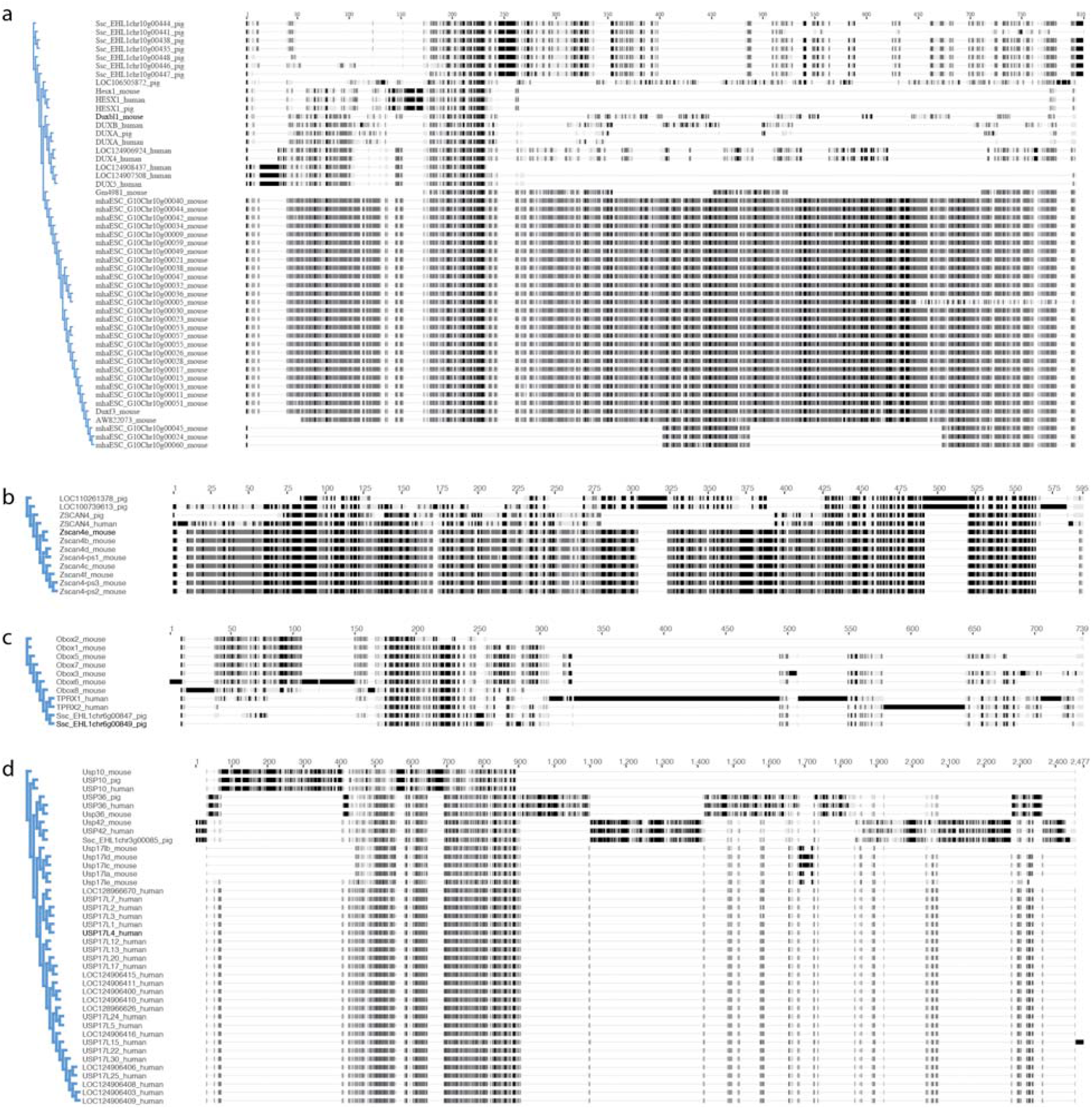
Protein-level alignments of representative totipotency genes (families) with divergent conservation patterns. Multiple-sequence alignment summaries for selected gene families, highlighting species-specific expansion and divergence. Panels show alignment profiles for **(a)** DUX-family proteins (including multiple pig DUX-like copies), **(b)** ZSCAN4 family, **(c)** OBOX family, and **(d)** USP17/USP17L family across pig, human, and mouse. The phylogenetic trees generated by neighbor-joining method are displayed on the left and horizontal tracks summarize aligned blocks and gaps across each protein.

**Supplementary Figure 11.**
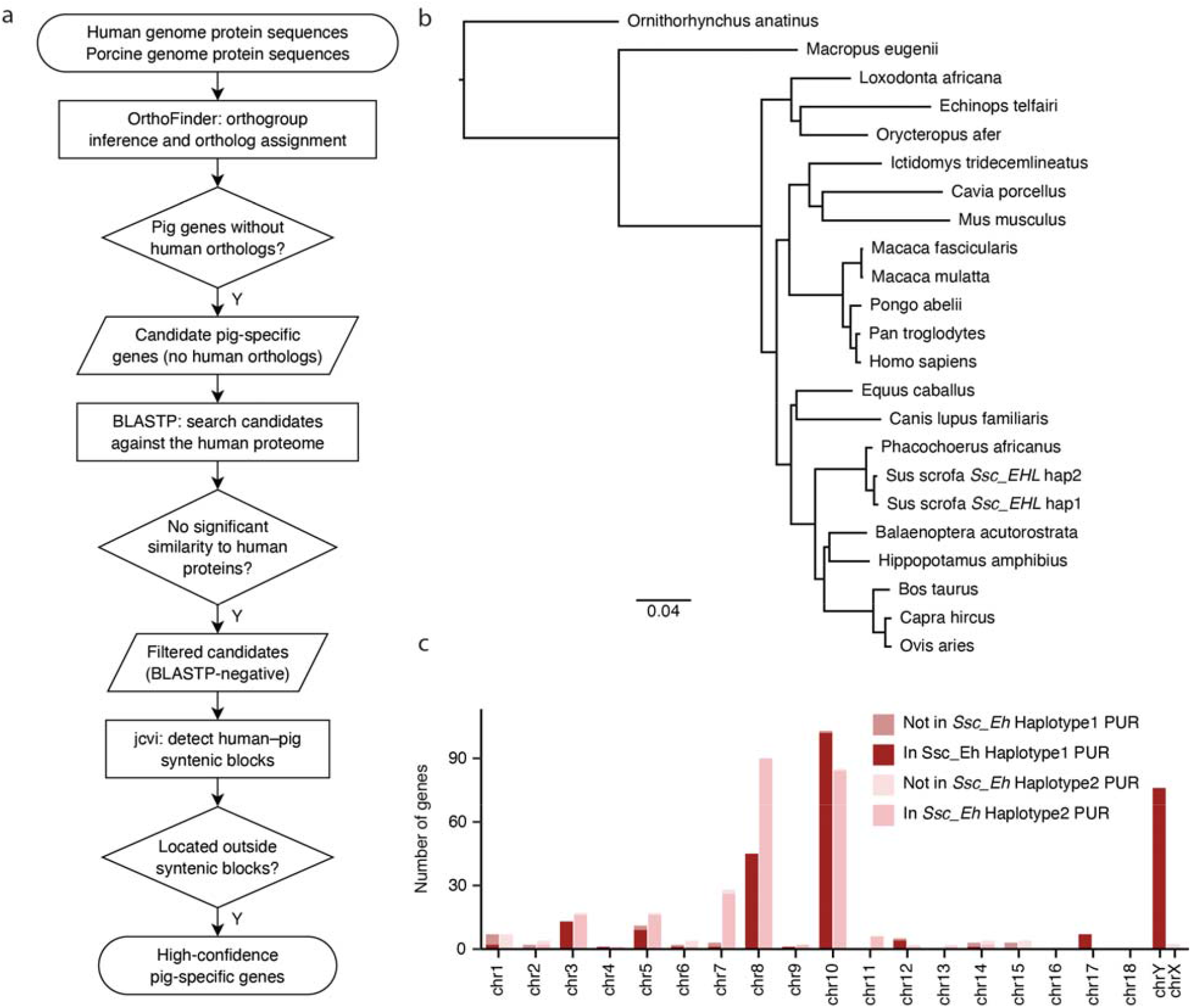
Definition of high-confidence pig-specific candidates and phylogenetic context for hit assignment. **(a)** Stepwise filtering pipeline for pig-specific candidate identification, integrating orthogroup inference, BLASTP screening against the human proteome, and synteny block detection to exclude candidates within conserved human-pig syntenic regions. **(b)** Phylogenetic relationships among representative mammals used to contextualize taxonomic assignment of top-scoring homology hits for pig-specific candidates (scale bar shown). **(c)** Chromosomal distribution of pig-specific genes, stratified by their presence within PURs.

**Supplementary Figure 12.**
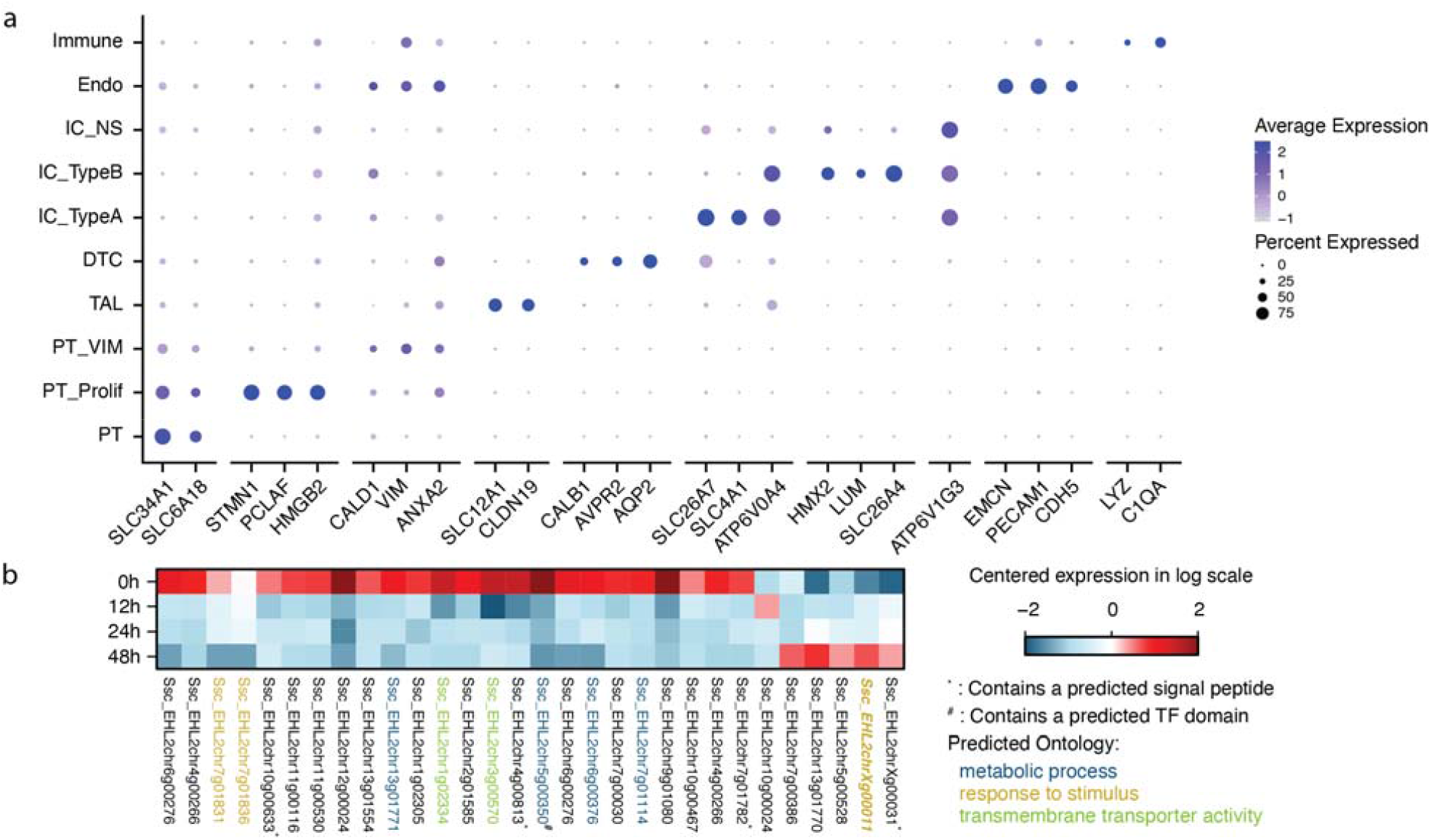
Reanalysis of xenograft transcriptomes and characterization of newly resolved pig genes in *Ssc_EHL*. **(a)** Dot plot showing canonical marker genes used to define major renal cell populations in the pig-to-human kidney xenograft single-cell RNA-seq dataset. Color intensity represents average gene expression, and dot size indicates the percentage of cells expressing each gene. Cell populations include immune cells, endothelial cells (Endo), nonspecific intercalated cells (IC_NS), type B intercalated cells (IC_TypeB), type A intercalated cells (IC_TypeA), distal tubule cells (DTC), thick ascending limb cells (TAL), proximal tubule cells (PT), vimentin-positive proximal tubule cells (PT_VIM), and proliferating proximal tubule cells (PT_Prolif). **(b)** Temporal expression patterns of newly resolved genes identified by reanalyzing longitudinal bulk RNA-seq data from pig-to-human kidney xenograft biopsies collected at 0, 12, 24, and 48 hours after transplantation. Gene expression values are shown as centered log-scaled expression. Symbols indicate genes containing a predicted signal peptide (^*^) or a predicted transcription factor domain (^#^). Gene labels are colored according to predicted functional categories, including metabolic process (blue), response to stimulus (yellow), and transmembrane transporter activity (green).

**Supplementary Figure 13.**
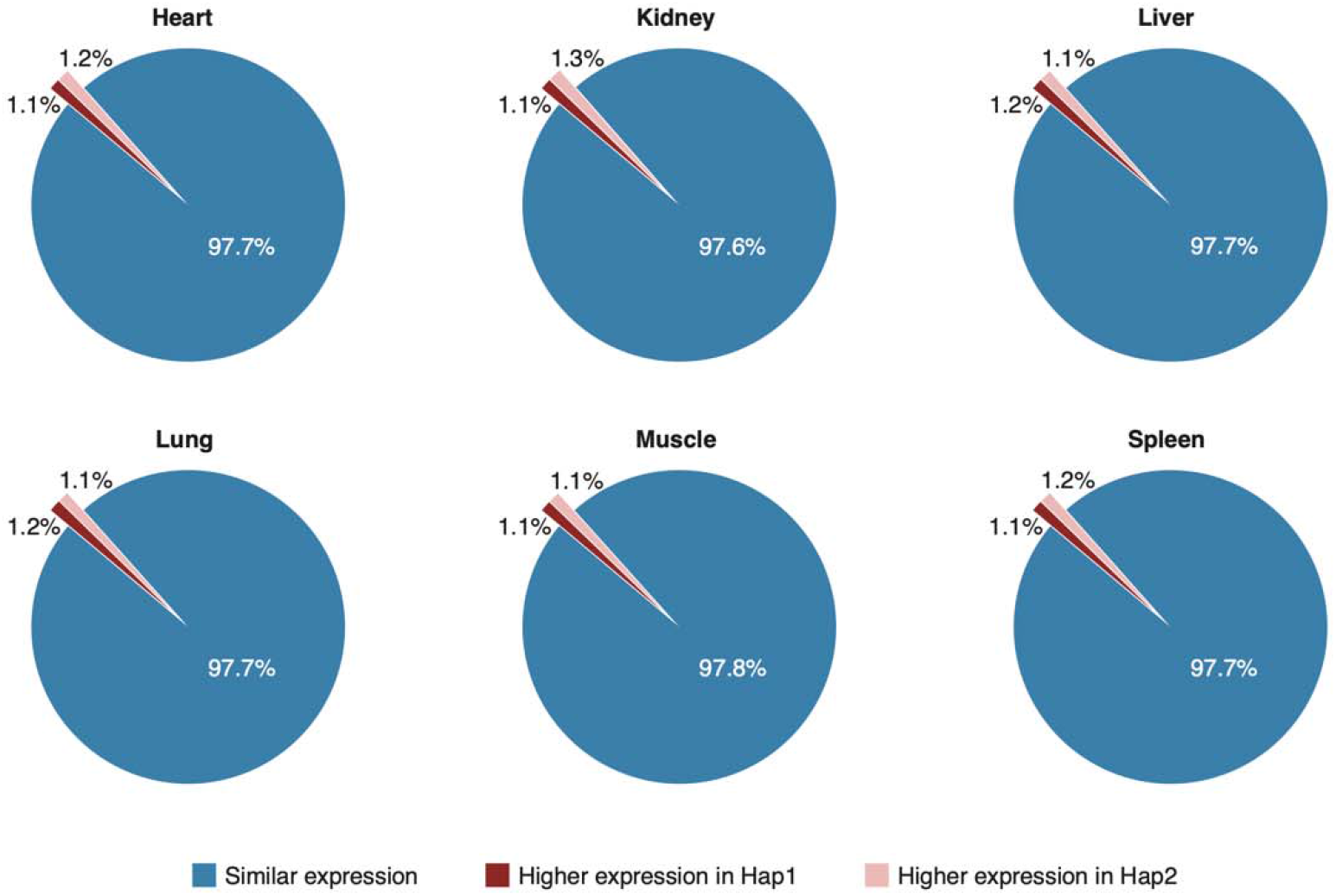
Haplotype concordance of tissue transcriptomes across six organs. Pie charts summarize the fraction of genes showing similar expression between haplotypes versus haplotype-biased expression in each tissue (heart, kidney, liver, lung, muscle, and spleen). Colors denote similar expression, higher expression in hap1, and higher expression in hap2.

**Supplementary Figure 14.**
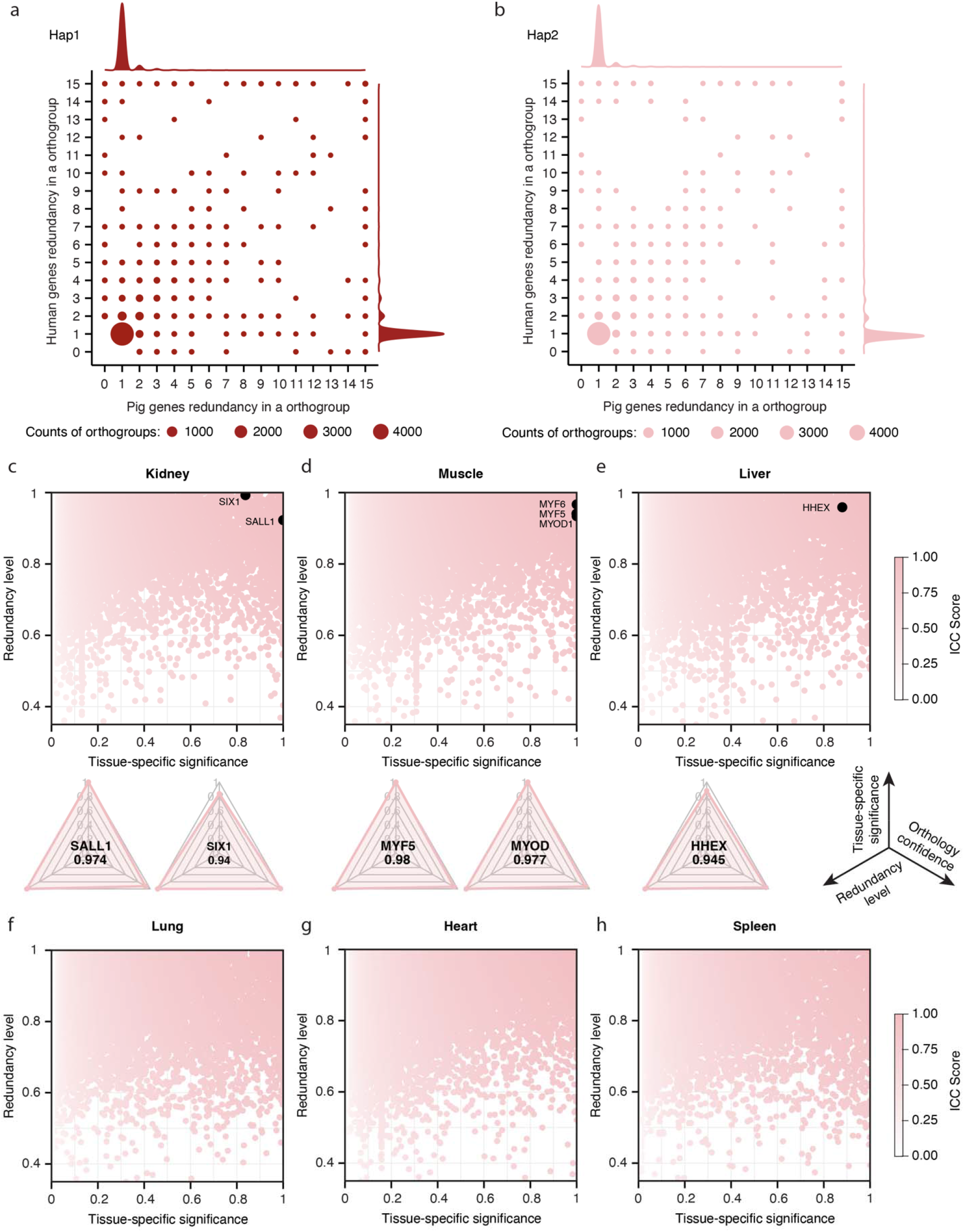
Organ-enriched gene sets and ICC landscapes on Ssc_EHL. **(a)** Pig-human orthogroup redundancy landscape shown for hap1-derived gene models. Points represent orthogroups plotted by pig versus human gene redundancy; point size denotes orthogroup counts. **(b)** Pig-human orthogroup redundancy landscape shown for hap2-derived gene models. **c-h**, ICC scores landscapes for kidney (**c**), muscle (**d**), liver (**e**), lung (**f**), heart (**g**), and spleen (**h**) on hap2, plotted as tissue-specific significance versus redundancy level with ICC score indicated by color. Previously reported interspecies chimerism target genes (e.g., *SALL1* and *SIX1* for kidney, *MYOD1, MYF5*, and *MYF6* for muscle, and *HHEX* for liver) are highlighted.

